# Motivation-dependent communication between cortex and brainstem

**DOI:** 10.64898/2026.11.05.764652

**Authors:** Timothy A. Machado, Yiming Chen, Manuel Beiran, Madhav Subramanian, Chandan Kadur, Susanna Bradbury, Kerry Nix, Brady Liu, Kanaka Rajan, Karl Deisseroth

## Abstract

Neurons in the medullary brainstem exhibit movement-related patterns of activity, but the neuronal population codes that specify distinct actions, and the mechanisms by which cortex may shape these codes as a function of motivational state, remain unclear. To explore whether coupling between cortex and medulla depends upon motivational context, we trained mice to make both pre-reward “appetitive” licks that served to trigger water availability and rapidly attenuated with satiety, and “consummatory” licks, which occur post-reward release and more slowly attenuated with satiety. Simultaneous recordings from secondary motor cortex and motor medulla revealed that medullary activity was more time-locked than cortical activity to individual licks and exhibited a phase-based population code that encoded lick direction. Correlations between cortex and medulla were stronger during appetitive licks than during consummatory licks, with causal dependency established by optogenetic intervention and supported by multi-region recurrent network models. Population codes underlying behavior shared across the central nervous system, exemplified by cortico-medullary coupling, thus may depend upon the motivational context underlying action.

---

Over the past two decades, many studies using head-restrained mice to study the cortical influence over movement have focused on licking behavior. In particular, a cortical subregion of the secondary motor cortex (MOs), known as anterolateral motor cortex (ALM), has been shown to be necessary for performing many licking tasks^1–3^. However, recent studies have demonstrated that silencing ALM does not in fact abolish all licking, but rather more selectively impairs specific features of licking movements^4–6^. These results suggest that, more generally, cortex may only exert modulatory influence over certain movements, perhaps restricted to certain motivational contexts^7,8^.

Moreover, in adult mice, the cortex provides little to no direct input onto motor neurons^9,10^. Instead, orofacial motor neurons, which reside in the brainstem, receive the majority of their direct inputs from nearby “premotor” interneuron circuits^11,12^. More specifically, both motor and premotor nuclei are contained in the medullary motor-related region, MY-mot (nomenclature from the Allen Brain Atlas taxonomy). Descending cortical neurons (arising mostly from primary [MOp] and secondary motor cortex [MOs]), in turn, provide much input to these premotor networks^13,14^. The ability of the cortex to control orofacial behavior therefore largely depends on its ability to affect the population activity of these circuits in MY-mot. However, owing to technical limitations, most earlier work has focused instead on directly relating cortical activity to movement—rather than cortical activity to premotor network activity. Therefore, relatively little is known about how cortical activity relates to that of the premotor network during any type of movement^3,13,15^.

Most knowledge of MY-mot function concerns its relationship with its motor outputs, rather than its long-range neuronal inputs. The MY-mot is known to contain a set of oscillator circuits that control rhythmic orofacial movements that are each entrained to the breathing cycle^16^. Along the dorsoventral axis of the medulla, there are oscillator circuits that control licking, chewing, whisking, and breathing. While most of these oscillatory circuits were identified decades ago^17–20^, we lack a complete understanding of how the circuits are organized into their constituent cell types and how they produce specific patterns of population activity that drive specific movements. *In vivo* recordings in the medulla have just recently become possible using modern tools for large-scale electrophysiology^3,13,21^, and optogenetics^19,22,23^. These studies revealed that there are multiple genetically-defined cell types distributed across the MY-mot that are tuned to whisking, licking, and breathing. But it still remains unclear how these orofacial oscillator circuits may encode and drive the detailed kinematics of individual orofacial movements.

One recent study measured neuronal population activity in the MY-mot during a directional licking task^13^. They identified neurons with firing modulated by licking, but the population code that relates MY-mot activity to specific lick trajectories (e.g. in different directions) was not determined. One idea comes from recordings from turtle spinal cord central pattern generating (CPG) circuits during reaching. This study identified populations of interneurons that are tuned to specific phases of a reaching action^24^. They found that each neuron fired at a specific phase during one type of reaching movement, but at distinct phases during a second type of reaching movement. This led us to wonder if a similar phenomenon may exist in the mammalian brainstem during distinct kinds of licking movements. If CPG circuits for licking in the mouse MY-mot were to function during intact behavior in a similar manner as the turtle spinal cord, the mean phase tunings of MY-mot neurons should systematically shift when the animal licks in different directions. In addition to demonstrating functional similarity between rodent brainstem CPGs and reptile spinal cord CPGs, this result would also highlight a specific temporal coding mechanism by which CPG circuits might represent distinct movements. Such phase codes may also be a key parameter that can be influenced by descending command pathways.

A second key question is how the neuronal circuits in the MY-mot integrate cortical input to generate behavior. Traditionally, MY-mot has been viewed as part of a rhythmic “motor plant” that produces movement-related patterns to drive motor neurons, operating under the influence of descending commands from cortex and other areas across the brain^25^. But the extent to which MOs and MOp can control the MY-mot, and whether the MY-mot merely propagates commands or instead actively interacts with cortical areas and encodes higher order contextual variables that could influence future actions, remains unclear. Analyses of large-scale electrophysiological recordings have identified neurons in MY-mot that encode non-motor information, such as future choice, which may be provided by cortical regions^26^. Neural recordings in a directed licking task showed that the influence of MOs and MOp during movement varied considerably over specific motor actions and over time, even over very short timescales, suggesting that the role of motor cortex during movement, and therefore its effect on the medulla, is dynamic^6^. Together, these recent findings suggest that MY-mot and motor cortex may be better understood as a system with bidirectional interactions that are dynamically adjusted on sub-second timescales. How such interactions are specific to certain motor actions, or during particular motivational contexts, remains unclear.

To address these questions, we first developed a head-fixed behavioral assay wherein mice make directed licks across variable behavioral contexts. During this task, we then performed population recordings in the MOs and across putative postsynaptic and premotor neuronal populations in the MY-mot. Analysis of our MY-mot data revealed a rhythmic phase-based population code for lick direction, and a canonical correlation-based decoder analysis revealed that consistent neuronal population codes for lick direction generalized even across distinct mice. Next, we analyzed data where we simultaneously recorded from across the MY-mot and MOs. These results support the idea that cortical control over licking behavior is dependent on motivational context; cortical influence is stronger during more dexterous “appetitive” licks and weaker during more rhythmic “consummatory” licks. Together, these results reveal principles by which the cortex and the brainstem flexibly work together to coordinate specific licking movements.

## RESULTS

### A lick-to-target task reveals a role for motivation across different lick types

We trained mice to perform a head-fixed behavioral task wherein they needed to make licks directed towards three different spouts **(Figure 1A, left),** in order to receive water rewards. Tongue movements were simultaneously captured using dual high-speed cameras and then reconstructed in 3D with a custom UNet-based segmentation pipeline (**Figure 1A, right**) similar to previous work^5^. All experiments were performed in VGAT-ChR2 mice, making it possible for us to transcranially inhibit MOs (by optogenetically stimulating inhibitory interneurons) as we concurrently performed electrophysiological recordings during natural licking behavior (**Materials and Methods**).

**Figure 1.**
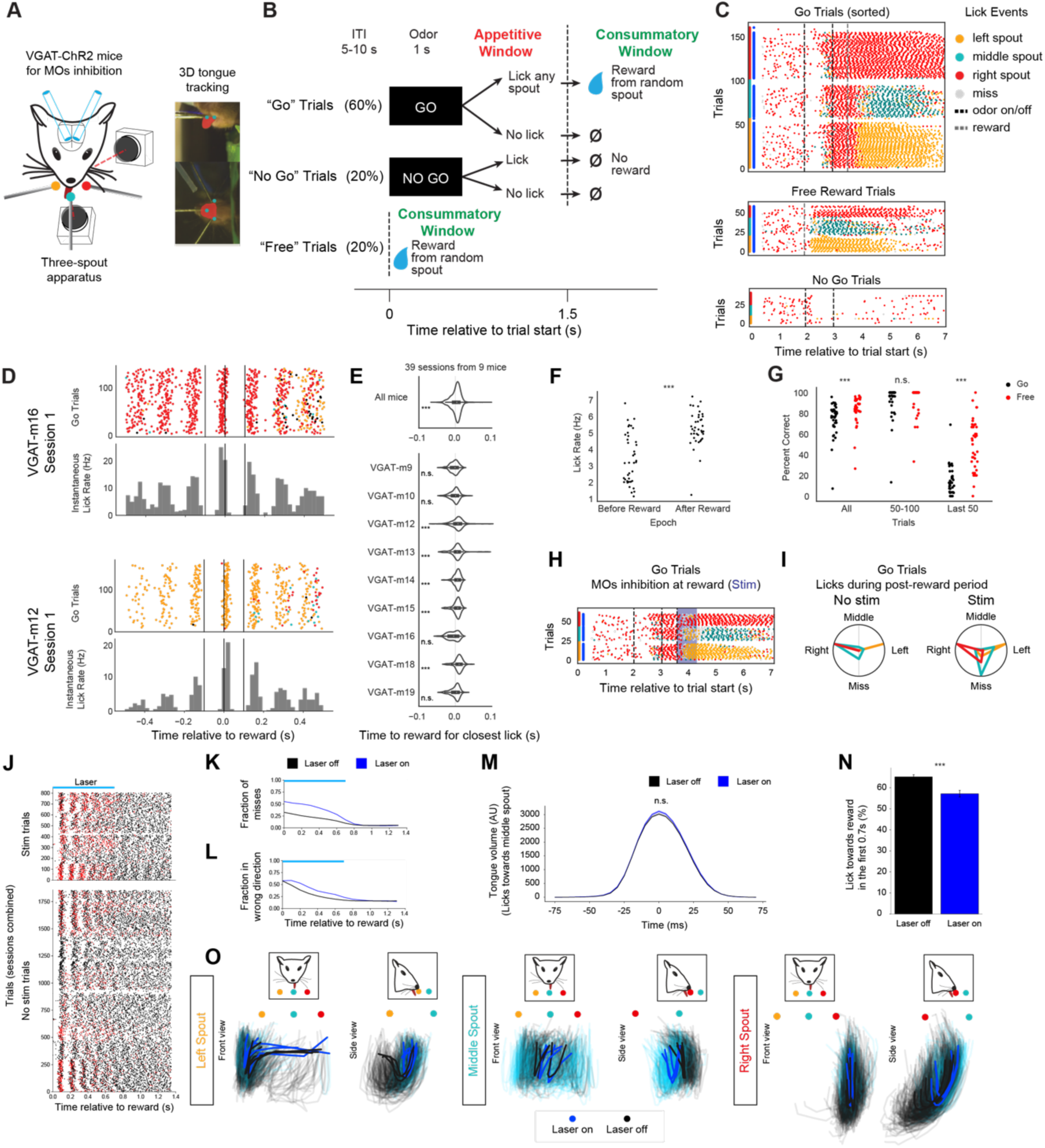
Go/no go directional licking assay reveals behavioral structure in goal-oriented licking. **A.** Left: Illustration of the three-spout experimental setup for directional licking behavior and optogenetics in VGAT-ChR2 mice. Right: Licking was quantified using two orthogonal high-speed cameras and a UNet model trained to segment the tongue from each view. **B.** Task structure for “go”, “no go” and “free” trials, indicating the appetitive and consummatory epochs of each trial type. **C.** Lick raster showing licks toward the left (yellow), middle (teal), right (red) spouts in addition to licks that missed a spout (gray) from a representative session (164 go trials; 64 free trials; 59 no go trials). Blue bars on the vertical axis indicate rewarded trials. Red, teal, and yellow bars on the vertical axis indicate the rewarded spout. Trials are sorted by type for visualization purposes. **D.** Lick raster plot and histogram with licks aligned to reward start, two representative sessions shown, with each row representing a trial (145 and 121 go trials, respectively; bin size = 0.02 s). **E.** Distribution showing the relative timing of licks that are closest to reward delivery (data from 9 mice with 160, 358, 306, 143, 144, 224, 180, 139 and 202 trials, respectively). A Wilcoxon test was used to test whether the median was greater than zero. **F.** Lick rate during appetitive and consummatory epochs of the task. **G.** Task performance (percent of correct trials) considering all the trials in each session, only trials 50-100 (mid-session), or only the last 50 trials. Each dot corresponds to a different session. **H.** Lick raster plot on “go” trials with optogenetic stimulation, for one example session. **I.** Distribution of different lick directions and missed licks in laser on and laser off trials. **J.** Lick raster plot showing missed (red) and hit (black) licks. **K.** Fraction of post-reward licks that miss the rewarded spout as a function of time after reward, in laser on (blue) and off (black) trials. **L.** Fraction of post-reward licks towards a non-rewarded spout as a function of time from reward, in laser on (blue) and off (black) trials. **M.** Average timecourse of the recorded tongue volume during licks towards the middle spout in laser on (blue) and off (black) trials. **N.** Percentage of licks towards the rewarded spout in the first 0.7 s after reward delivery, in laser on (blue) and off (black) trials. **O.** Front and side view of the tongue tip trajectories, grouped by lick direction. Trajectories recorded during the laser on period are shown in blue; and trajectories recorded during the same time window in laser off trials are in black. (ns: p >= 0.05; *: 0.05 > p >= 0.01; **: 0.01 > p >= 0.001; ***: p < 0.001)

The behavioral task consisted of three different trial types (**Figure 1B**). On “go” trials, in response to an olfactory cue, mice were required to first make “appetitive” licks towards any water spout in order to trigger the release of water from a random water spout that they would then consume (by making “consummatory” licks). Mice often had to change lick directions in the middle of a rhythmic bout, when the chosen spout during appetitive licking differed from the spout that was randomly chosen to release water reward on that trial. In contrast, on “free” trials, mice received water from a random spout without any prompting stimulus. Importantly, the spouts were positioned in contact with the mouse’s face such that water rewards could be felt by the mouse when they were released. On “no go” trials, mice learned to withhold licking in response to a “no go” associated odor cue as they were not rewarded regardless of their behavior.

We found that after two to three weeks of training, mice could reliably perform this task, receiving reward on more than 80% of “go” trials in the middle of a session (**Figure 1C and Figure S1D**). Moreover, trained mice also exhibited highly stereotyped licking behavior both in the timing of individual licks, and in their idiosyncratic preferences for which spouts to lick during the “go” odor cue. We used a custom 3D computer vision pipeline to track the time of maximum tongue protrusion with high temporal fidelity at 200 Hz (**Materials and Methods**). Analysis of tongue protrusion events revealed that mice learned to precisely synchronize the timing of their licking relative to the timing of expected reward on “go” trials (**Figure 1D-E**; the lick rate is defined as the mean number of licks per bin across trials, multiplied by the 0.02 s bin width). We found that, after training, most mice consistently timed their licking such that their tongues were already being protruded prior to reward release so that they would make spout contact precisely around the time of reward delivery (median lick–reward offset across mice = 1.7 ± 0.3 ms; n=1,854 licks from 9 mice), with a significant positive median in 6/9 mice (Wilcoxon signed-rank test vs 0, p<0.001).

Further analysis of licking on “go” trials revealed that mice consistently increased their rates of licking upon the onset of reward. This licking rate difference thus represents a behavioral distinction between pre-reward “appetitive” licks and post-reward “consummatory” licks (p<0.001, Wilcoxon signed-rank test, n=39 sessions from 9 mice, including licks within 0.5 s of reward, excluding the lick closest to reward, on correct go trials with no optogenetic stimulation, **Figure 1F**). We also found that mice were significantly less successful at obtaining rewards on “go” trials than on “free” trials (p<0.001, Wilcoxon signed-rank test, comparing trial success rate between all “go” and “free” trials; n=39 sessions from 9 mice, **Figure 1G, left**). Importantly, performance did not differ significantly between these two trial types in the middle of each session, but as the mice became sated during later trials, they were less likely to make “appetitive” licks to trigger reward release while continuing to make “consummatory” licks in response to free rewards that they could feel on their face (p>0.05, Wilcoxon signed-rank test, comparing success rate between “go” and “free” trials during middle session trials 50-100; p<0.001, during the last 50 trials of each session; n=39 sessions from 9 mice, **Figure 1G, middle and right**). This result indicates that “appetitive” licks require higher levels of motivation than “consummatory” licks, and therefore might be controlled by different neuronal mechanisms.

Previous studies have implicated MOs in diverse aspects of licking control, ranging from necessity for lick generation to fine modulation of tongue trajectories^1,5,27^. To determine how MOs may contribute to licking within our task, we bilaterally inhibited MOs at the time of reward delivery (**Figure 1A, Materials and Methods**). Consistent with prior reports ^5^, optogenetic inhibition increased the rate of licks that “miss” (defined as licks in which the tongue did not reach the rewarded spout, **Figure 1H-L**). We then investigated what kind of error caused these misses. First, we observed no systematic nor significant change in overall lick rate or in gross tongue volume (a proxy for tongue protrusion; **Figure 1M**; AUC: no stimulation 20,561 ± 291 (n=1,368 licks) and during stimulation 21,195 ± 360 (n=945 licks)). Next, during MOs inhibition, we detected a decreased fraction of licks that were directed towards the rewarded spout (**Figure 1L, N and Figure S1A-C**; ratio towards reward in no stimulation trials: 0.65 ± 0.01 (n=1,233) and during stimulation 0.57 ± 0.02 (n=508), p<0.001 Mann–Whitney U test). Finally, tracking of tongue tip trajectories revealed specific disruptions in licking kinematics relative to the target spouts (**Figure 1O and Figure S1B**). In summary, we found that bilateral inhibition of MOs at the onset of reward did not cause a gross disruption of licking behavior. Instead, it increased the fraction of errant licks towards unrewarded spouts, and caused mice to make short, hypometric licks. This is most apparent with licks towards the most-distant middle spout (**Figure 1O, middle**).

**Figure S1.**
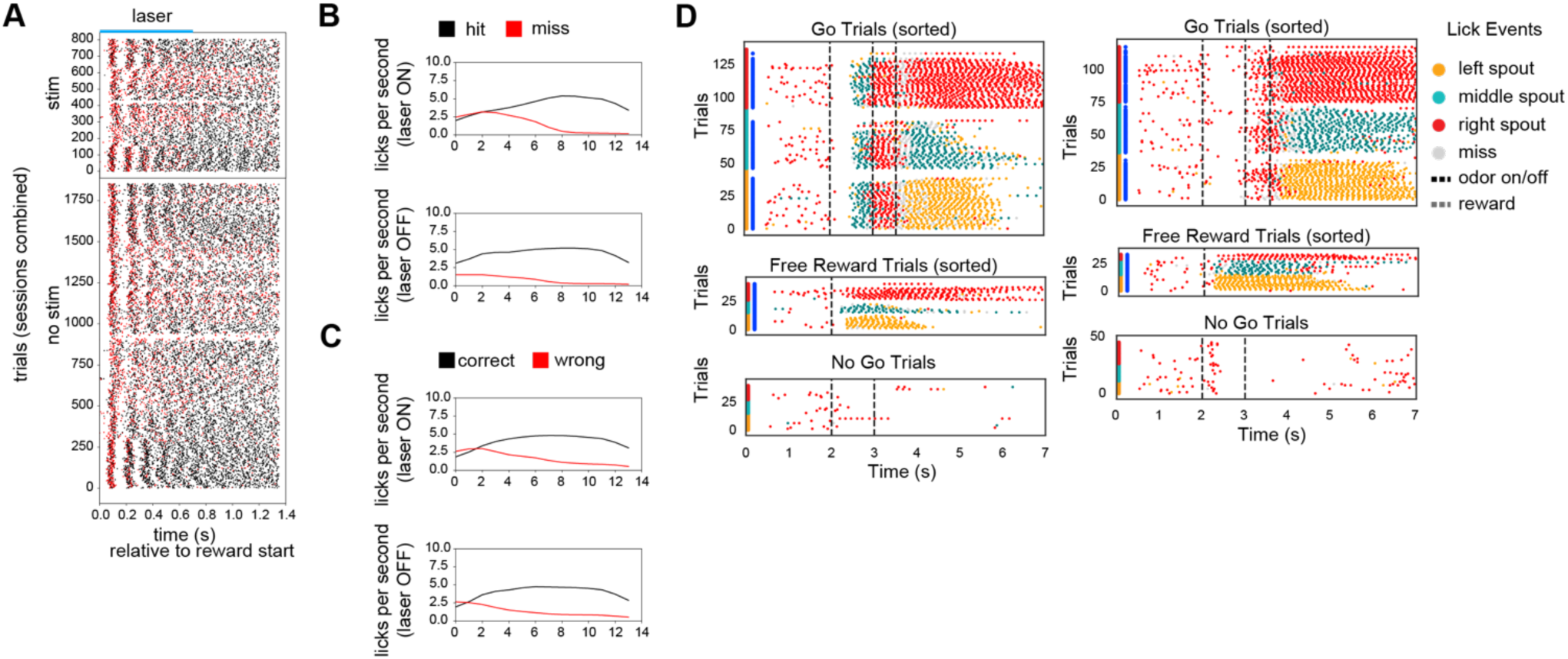
Go/no go directional licking assay reveals behavioral structure in goal-oriented licking. **Relates to Figure 1. A.** Representative lick raster plot showing wrong (red) and correct (black) licks from a single session in a VGAT-ChR2 mouse. Licks are considered as wrong when they target a non-rewarded spout. **B.** Rate of missed (red) and hit (black) licks vs time in laser on (top) and off (bottom) trials. **C.** Rate of wrong (red) and correct (black) licks vs time in laser on (top) and off (bottom) trials. **D.** Lick raster showing licks toward the left (yellow), middle (teal), right (red) spouts in addition to licks that missed a spout (gray) from two representative sessions with different mice. Blue bars on the vertical axis indicate rewarded trials. Red, teal, and yellow bars on the vertical axis indicate the rewarded spout. Trials are sorted by type in these plots for visualization purposes.

### Encoding of task and tongue kinematics in cortex and medulla

Neurons in both the secondary motor cortex (MOs) and medullary motor nuclei (MY-mot) have been implicated in the control of licking and orofacial movements^1,2,28^. To understand how licks are encoded by putative premotor circuits in the brainstem, we recorded units in cortex and/or medulla in each animal during our behavior task, and post-hoc registered each probe track to the Allen brain atlas (**Figure 2A-C**)^29^. Consistent with previous work^4,5,13^, the majority of units recorded in our task exhibited temporally structured firing patterns aligned to distinct stages of the licking sequence (**Figure 2C–E**; in D: 818 MOs units and 509 MY-mot units). Despite the distinct anatomical and functional identities of MOs and MY-mot, principal component analysis (PCA) and unsupervised clustering of trial-aligned population activity revealed qualitatively similar principal components and k-means clusters in both regions (**Figure 2D-E and Figure S2A-B**; variance explained by first 4 PCs: 55.5% in MOs dataset and 60.9% in MY-mot), suggesting convergent representations across task epochs.

**Figure 2.**
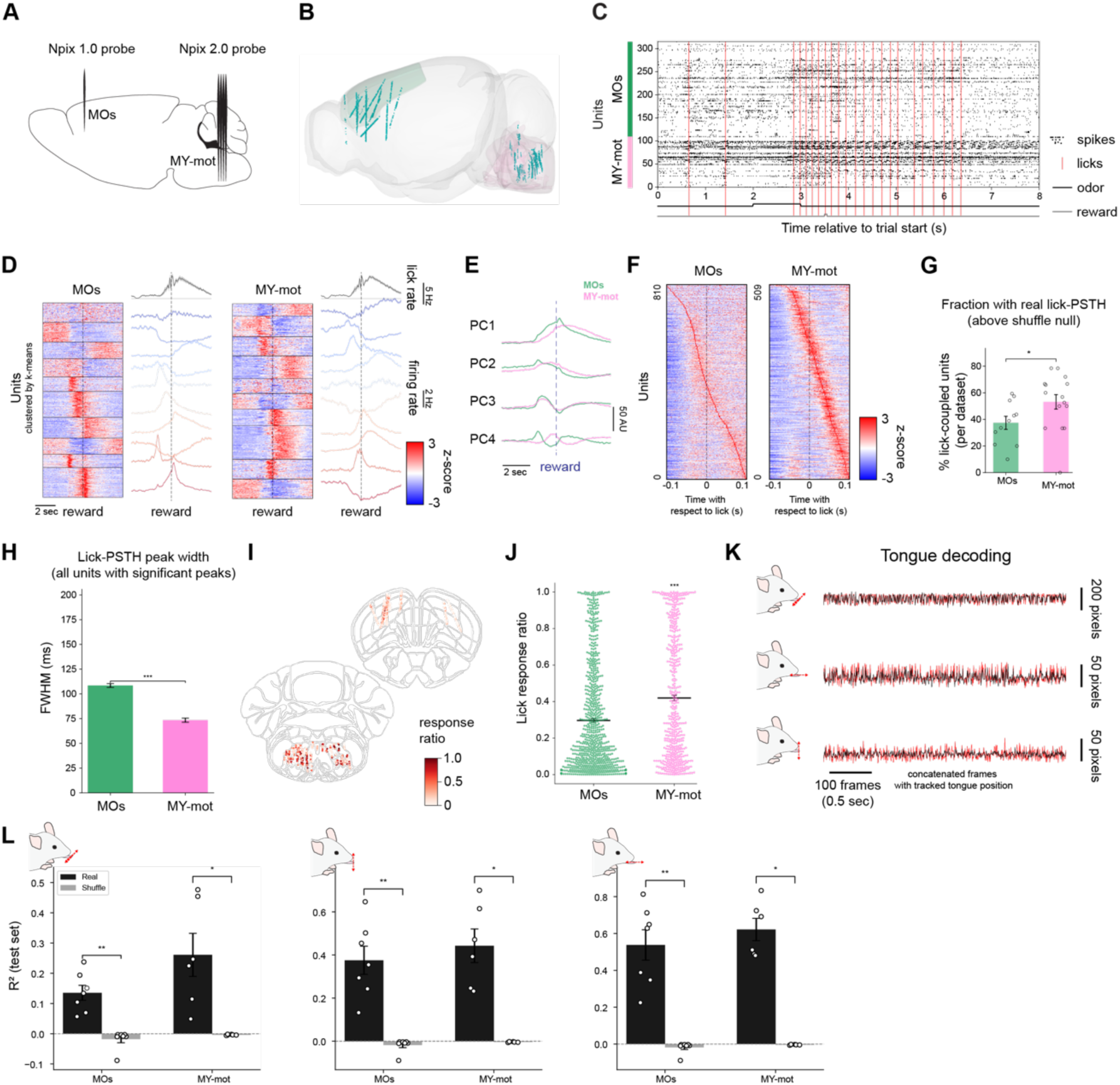
Cortical and medullary units are correlated with licking behavior and encode tongue kinematics. **A.** Illustration of the dual Neuropixels recording setup. **B.** Reconstructed location of recorded units, aligned to the 3D atlas. **C.** Spike raster plots of MY-mot and MOs activity in one example ‘go’ trial. **D.** Peri-stimulus time histogram (PSTH) of activity from MOs and MY-mot units aligned to reward delivery (dashed vertical line). Units were clustered using k-means (in each cluster: MOs N=82, 83, 69, 77, 61, 109, 87, 68, 56 and 126; MY-mot N=32, 54, 50, 59, 39, 41, 43, 88, 51 and 52). Average firing rate and SEM (shaded) for each cluster are shown to the right side of each panel. **E.** First four PCs of MOs and MY-mot activity during ‘go’ trials. **F.** Peri-lick histogram of MOs and MY-mot units: normalized firing rate averaged across licks and aligned to the time of maximal protrusion (MOs N=810; MY-mot N=509). **G.** Percentage of units per dataset whose lick-PSTH peak exceeded the 95th percentile of a circular-shift shuffle null. **H.** Full width at half maximum (FWHM) of peri-lick histogram of MOs and MY-mot units. **I.** Location of units across all experiments, on top of a 2D atlas map. Colors indicate the single unit response ratio: the fraction of licks to which each unit spikes. **J.** Response ratio (ratio of licks with at least one spike) of units recorded from MOs (N=821; 0.297±0.01) and MY-mot (N=509; 0.419±0.015). **K.** Linear regression predictions of 3D tongue tip positions based on the population firing rate (black: recorded position; red: predicted position). The model was trained with MY-mot data. **L.** R² of linear regression predicting tongue position (X, Y, Z axes) from population firing rates for MOs (n = 7 datasets) and MY-mot (n = 6 datasets), compared to a shuffle null in which the time axis of firing rates was permuted (200 permutations). (ns: p >= 0.05; *: 0.05 > p >= 0.01; **: 0.01 > p >= 0.001; ***: p < 0.001).

When neural activity was aligned to individual lick events, MY-mot units exhibited markedly tighter temporal coupling to single licks compared to MOs units (**Figure 2F-H and Figure S2C-I**). Correspondingly, single units in MY-mot units fired much more consistently during all licks than single units in MOs (**Figure 2I–J**; response ratio [the fraction of licks in which a unit fires at least one spike]: 0.297 ± 0.010 (n=821) for MOs units and 0.419 ± 0.015 (n=509) for MY-mot units, p<0.001 Mann–Whitney U test). This is consistent with the known role of medullary circuits in fine motor execution and timing^25^.

To test whether neural populations encode precise tongue kinematics, we trained linear-regression models on either MY-mot or MOs population activity to predict the 3D coordinates of the tongue tip trajectory. Models predicted tongue tip position consistently in all animals throughout trials in both MOs and MY-mot datasets (**Figure 2K-L; Figure S2J-L**).

**Figure S2.**
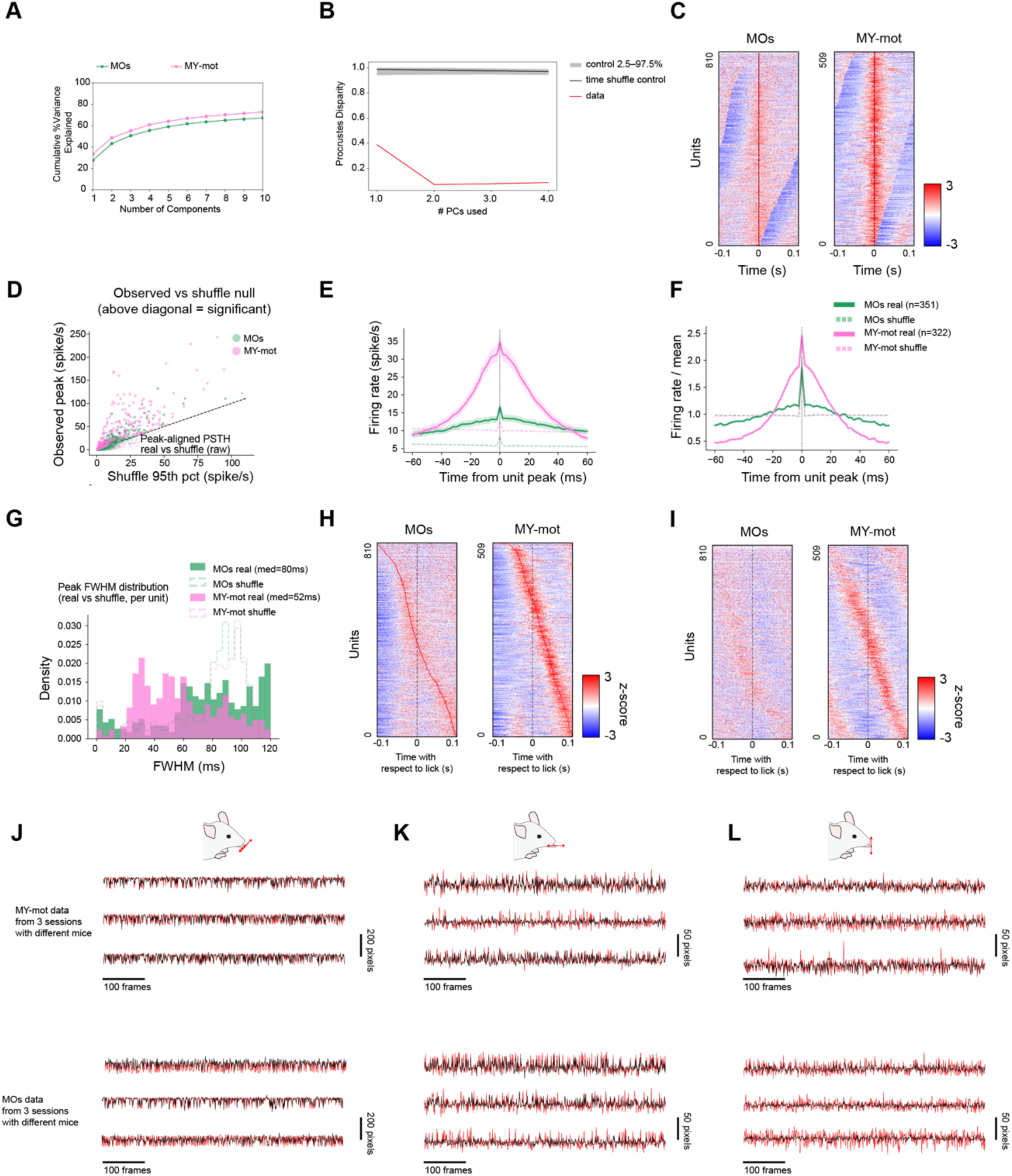
Cortical and medullary units are correlated with licking behavior and encodes tongue kinematics. **Relates to Figure 2. A.** Cumulative variance explained as a function of the number of principal components. **B.** Procrustes disparity between MY-mot and MOs subspaces as a function of the number of principal components. **C.** Peri-lick histogram of MOs and MY-mot units circularly permuted so that peaks are centered (MOs N=810; MY-mot N=509). **D.** Observed peak amplitude vs shuffle 95th percentile per unit; dashed line is the identity line, points above are significant. **E.** Mean-normalized peak-aligned PSTHs for significant units in MOs and MY-mot (real, solid; circular-shift shuffle, dashed). Shading shows SEM. **F.** MOs vs MY-mot real-data overlay showing regional difference in peak width. **G.** FWHM distributions per region. **H-I**. PSTH sorted by peak time of firing rate in even-lick PSTHs (left), with that order applied to odd-lick PSTHs (right). **J-L.** Linear regression model using firing rates to predict concatenated lateral (**J**), anterior-posterior (**K**) and vertical (**L**) tongue tip positions from across 3 sessions with no-lick intervals removed (black: recorded position; red: predicted position).

### A population phase code in the medulla for lick direction

Timing of spikes can control rhythmic movements, and thus we next asked whether spiking activity carries a phase code for lick direction^24^. Measuring the instantaneous lick phase of each spike relative to the times of max tongue protrusion revealed that many units, particularly those in MY-mot, were tuned to specific phases of the licking cycle (**Figure 3A**), consistent with phase-coding mechanisms in rhythmic motor systems^30^. Preferred phases tiled the entire lick period (**Figure 3B**), but phase tuning was stronger in MY-mot than in MOs (**Figure 3C–D**; Statistics from Rayleigh test on all spikes from each unit: vector strength : 0.053 ± 0.004 (n=808) for MOs units and 0.200 ± 0.010 (n=509) for MY-mot units, p<0.001 Mann–Whitney U test).

**Figure 3.**
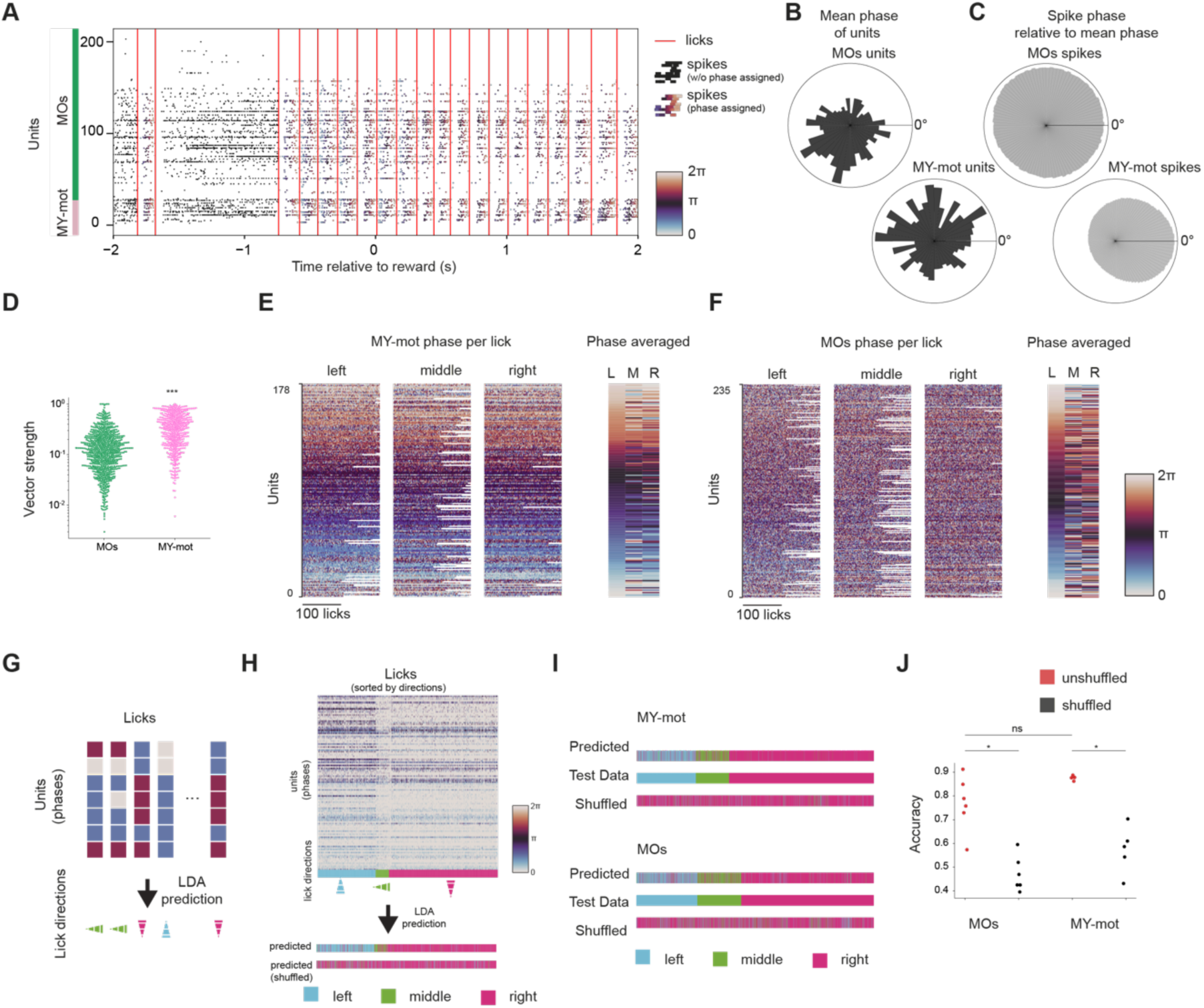
Temporal information of spikes encodes lick direction. **A.** Spike raster plot of an example trial, where the color of each spike represents their assigned phase. **B.** Preferred phase distribution across all units recorded from MY-mot and MOs. Zero degrees corresponds to the maximum tongue protrusion. **C.** Distribution of relative phases across spikes recorded from MOs and MY-mot units. The relative phase is defined as the difference between the phase of individual licks and the preferred phase of the spiking unit. **D.** Vector strength across units. MOs (N=808; 0.053±0.004), MY-mot (N=509; avg 0.200±0.010). **E.** Phase of spikes with respect to the licking cycle of MY-mot units. Licks are grouped by lick direction, and units have been sorted by preferred phase. **F.** Phase of spikes of MOs units with respect to each lick cycle. **G.** Illustration of the prediction of lick direction based on the phases of spiking activity using linear discrimination analysis (LDA). **H.** LDA predicted lick directions using phase and vector data in one example dataset, using both MY-mot and MOs phases. **I.** LDA prediction results when only MY-mot or MOs units were used. **J.** Accuracy of predictions (percentage of correctly predicted lick directions) across all datasets. (ns: p >= 0.05; *: 0.05 > p >= 0.01; **: 0.01 > p >= 0.001; ***: p < 0.001).

We then asked how the phase tuning of single units depends on the licking direction. We computed the circular mean phase value for each lick direction across units. We observed changes in the preferred phase based on lick direction (**Figure 3E-F, Figure S3**; Pairwise difference of average phase: 3.314 ± 0.117 (n=245) and 1.919 ± 0.090 (shuffled, n=392) in MOs, 1.628 ± 0.118 (n=178) and 0.525 ± 0.043 (shuffled, n=330) in MY-mot, p<0.001 Mann–Whitney U test). Based on this observation, we hypothesized that phase information from both cortex and the medulla may be sufficient to encode lick direction. To test this, we trained linear discriminant analysis (LDA) models using only the phase of the spike timing across units to predict lick direction (**Figure 3G-H**). Indeed, we found that both MY-mot and MOs populations could predict lick direction significantly above chance (**Figure 3I-J, Figure S3J-L**; average prediction accuracy: 0.77 for MOs, 0.47 for MOs shuffled, p<0.05 Mann–Whitney U test; 0.88 for MY-mot and 0.57 for MY-mot shuffled, p<0.05 Mann–Whitney U test;). These results indicate that both cortical and brainstem circuits carry temporally precise, phase-locked representations of tongue movements, with MY-mot exhibiting stronger temporal coherence and directional tuning.

**Figure S3.**
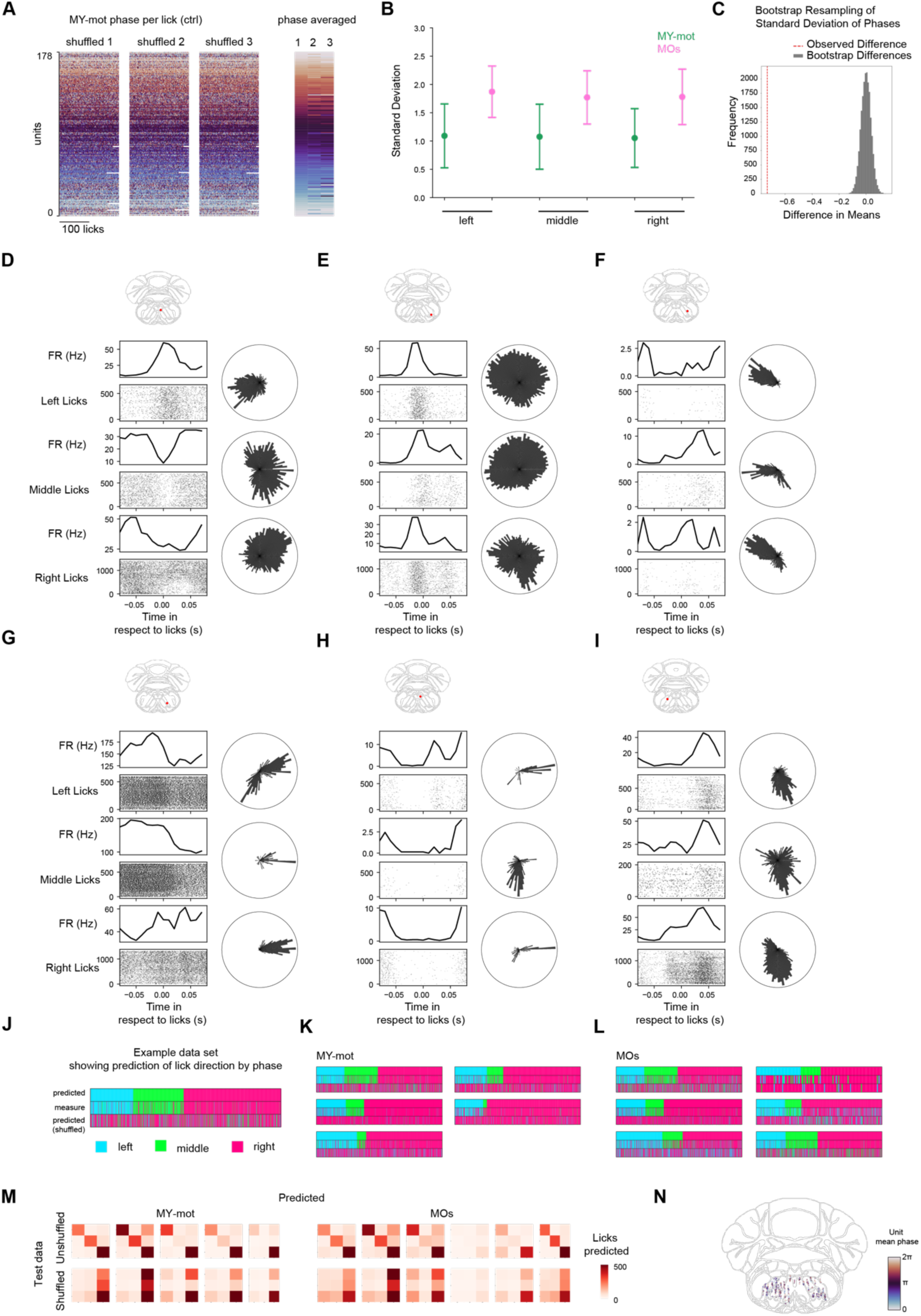
Temporal information of spikes encodes lick direction. **Relates to Figure 3. A.** MY-mot units (each row) and their phase coupling to each lick (each column), licks are sorted randomly. **B.** Bootstrapped standard deviation of phases. **C.** Bootstrapped distribution of estimates obtained from 10,000 resamples. The red vertical line marks the bootstrap mean. Histogram bars indicate the empirical distribution across resamples. **D-I.** Example phasic units from MY-mot, showing differential phasic response to licks towards different directions. **J-L.** LDA prediction results of individual MY-mot (**K**) and MOs (**L**) datasets combined. **J** illustrates how data are presented. **M**. Confusion matrices from LDA classification. Rows indicate true lick classes and columns indicate predicted classes. **N.** MY-mot units, color-coded by their mean lick phase, overlaid on a representative coronal atlas section. The medium section of all recorded slice is chosen to present.

### A shared population code for rewarded lick direction across subjects

Thus far, we have demonstrated that individual cortical and medullary units pooled from all recorded animals and sessions encode features of tongue kinematics and exhibit temporal firing patterns that vary with lick direction. This analysis assumes that there is a conserved neural representation of the task across individuals of the same species^31^. However, rather than presupposing that the shared code must take the form of phase, we asked a more general question: is there a consistent neural representation of rewarded lick direction across animals, regardless of the specific coding scheme? Whether such shared representations are present in the medulla is currently unknown^31^.

To address this, we applied multiset canonical correlation analysis (CCA) to dimensionality-reduced (PCA) activity from medullary premotor networks recorded in five mice while they performed the licking task described above (**Figure S4A)**^32^. CCA identifies shared latent dimensions by maximizing the variance of a consensus representation across datasets, yielding trajectories for each mouse in a common latent space (**Figure 4A**). The variance explained by aligned canonical components ranged from 6-16% in “free” trials and 13-23% in “go” trials (**Figure S4B-C**). Within this shared space, medullary population activity began to diverge shortly after mice began to lick towards distinct rewarded spouts (**Figure 4B**). The timing of this divergence tracked the increase in correctly directed licks, suggesting that the aligned latent dynamics reflected task-relevant structure and that the observed latency reflected the time interval when mice were still in the process of orienting their licks towards the location of the reward.

**Figure 4.**
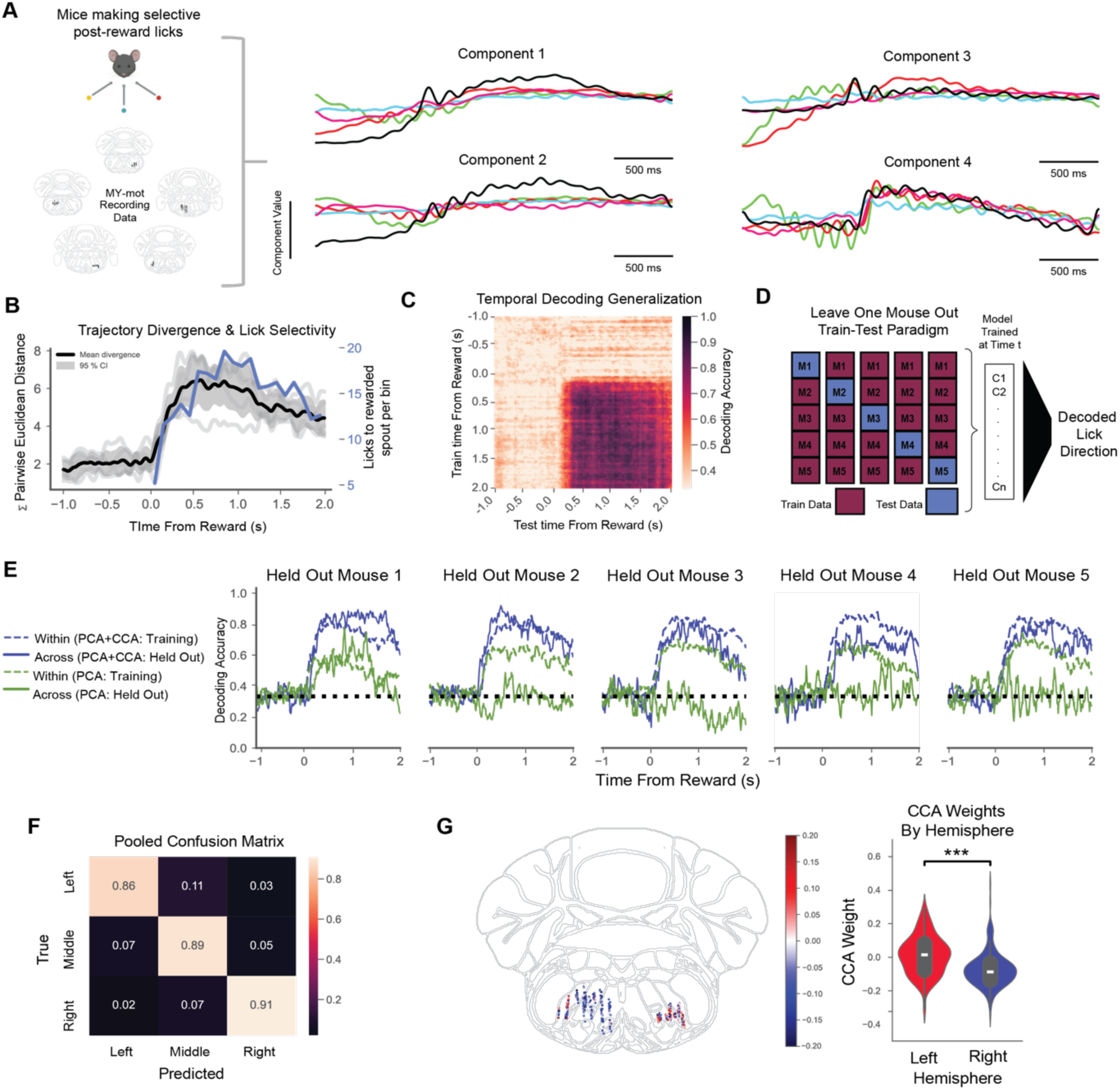
A shared medullary code that encodes lick direction. **A.** Schematic for aligning latent space. Principal components from each mouse were aligned using multiset canonical correlation analysis (CCA) to identify shared subspaces across mice. Trial-averaged trajectories for the first four canonical components are shown, with each color representing one mouse. **B.** Mean divergence of canonical trajectories across rewarded lick directions over time relative to reward (black line). The black trace shows the mean across 5 animals (±95% CI, shaded). Gray lines depict traces from individual mice. The blue trace indicates the number of licks directed toward the rewarded spout per time bin: mean across 5 mice. **C.** Temporal decoding generalization: classifiers trained at each time bin (–1 to 2 s) and tested across all bins show sustained cross-temporal decoding of rewarded lick direction, indicating stability in the medullary representation. **D.** Schematic for leave-one-mouse-out training and testing paradigm. At each time bin within the trial window, we trained a multinomial logistic regression classifier using data from four mice. Training included 5-fold cross-validation: models were fit on four folds and validated on the remaining fold. To assess cross-subject generalizability, the trained model was then evaluated on an entirely held-out mouse not included in training. This procedure was repeated such that each mouse served as the held-out test set once, ensuring the results reflect performance generalizable across all datasets. **E.** Comparison of decoding accuracy performance between aligned (CCA) and unaligned (PCA) data demonstrates that CCA alignment enables cross mouse decoding. **F.** Pooled confusion matrix of decoding accuracy (during 0-2s time window) in held out mice. **G.** Back projection of CCA weights highlights a dimension that encodes laterality in MY-mot units. Wilcoxon rank-sum test was used to calculate p-value. ***: p < 0.001.

The increase in trajectory divergence at the onset of consummatory licking, reflecting the task structure, suggests that the shared latent space derived from the five mice may encode the rewarded lick direction. To test this, multinomial logistic regression classifiers were trained at each time bin and tested across all bins. Accuracy was close to chance before reward delivery but improved at later time points (**Figure 4C**). Notably, classifier performance remained high even when applied to post-reward time bins not used in training, indicating that the aligned trajectories carried a relatively stable signal for rewarded lick direction (**Figure 4C, Figure S4D**).

Given that the aligned latent spaces encode sufficient information to enable decoding of lick direction, we wanted to test if these models could make predictions across mice. To assess cross-animal generalizability, we implemented a leave-one-mouse-out paradigm. Classifiers trained on four mice and validated on both held-out folds and held-out animals achieved comparable performance when data were aligned with CCA (**Figure 4D-E**). In contrast, unaligned PCA data maintained performance within folds but showed little generalization across animals. Confusion matrices summarized trial-level decoding performance in the aligned condition (**Figure 4F, Figure S4E**). We then asked whether medullary motor and premotor areas encode rewarded lick direction differently. We aligned neuronal recording from the hypoglossal nucleus (i.e. putative tongue-innervating motor neurons) to the consensus latent space of the medullary premotor network. Models trained on data from the medullary premotor network failed to successfully decode reward lick direction from hypoglossal nucleus neural activity (**Figure S4F**).

Earlier work demonstrated the presence of shared latent dynamics in the motor cortex during a reaching task^31^. By applying CCA to cortical recordings across individuals, we also found that alignment of neuronal activity data from the MOs can reliably enable superior cross-animal decoding of rewarded lick direction in comparison to unaligned data (**Figure S4G**). Decoding accuracy is comparable to aligned medullary data (**Figure S4H**).

The ability of a series of linear transformations (PCA and CCA) to align and enable cross animal decoding of rewarded lick direction is particularly surprising given that medullary data was recorded bilaterally (**Figure 2I**). To identify how the shared space accounted for data collected from different hemispheres, we projected CCA weights of the canonical component that explained the most amount of variance onto units (**Figure S4B and Figure S4I**). In doing so, we uncovered that a simple sign flip in weights helped align neural data from different hemispheres to enable decoding across hemispheres and animals (**Figure 4G**). In contrast, PCA weights showed no comparable laterality bias, suggesting that this feature emerged specifically from the canonical alignment (**Figure S4J**). Together, we have demonstrated a simple method to align neuronal data across hemispheres and animals, revealing a shared medullary premotor code that encodes reward lick direction.

**Figure S4.**
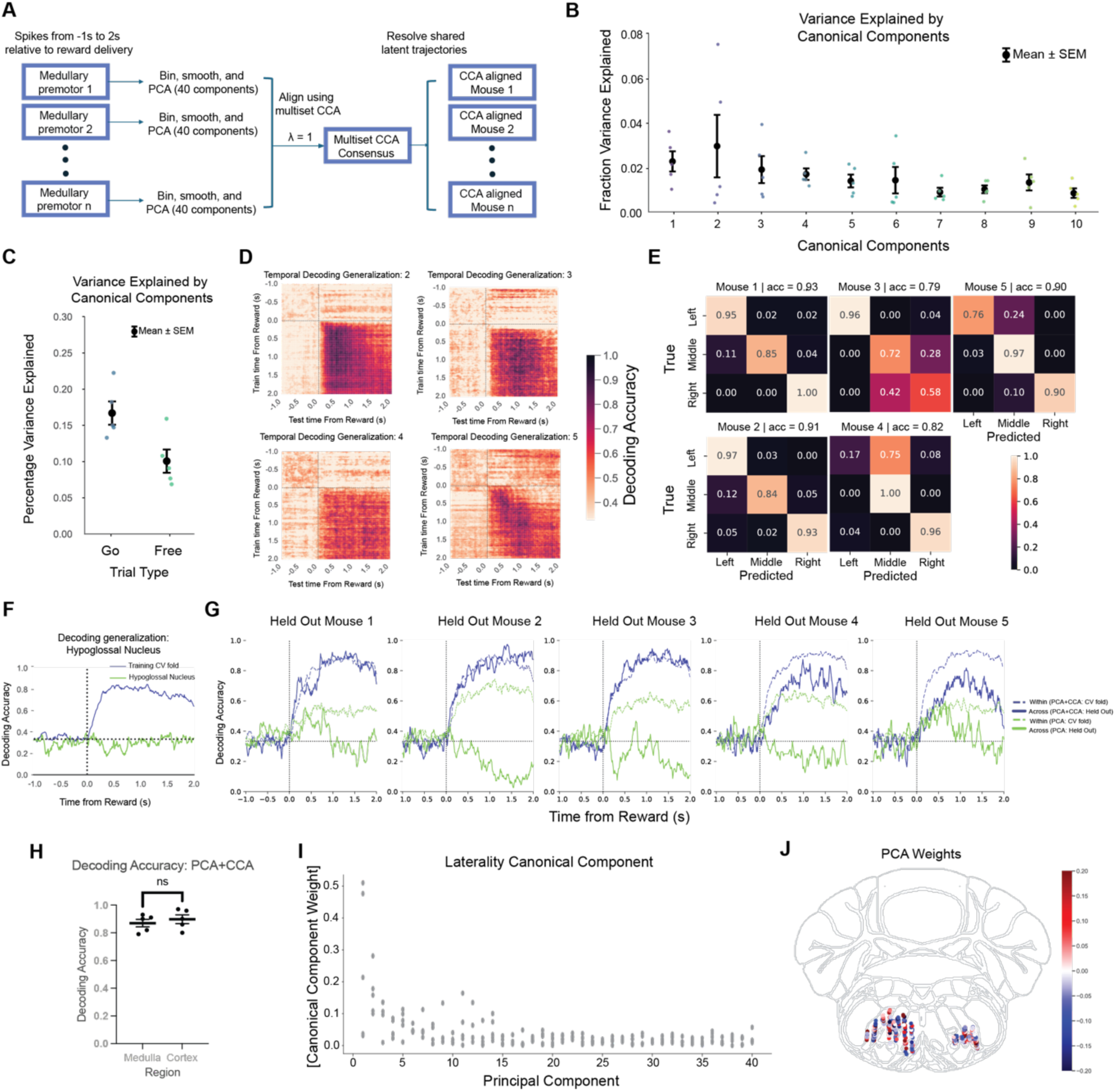
Multiset CCA identifies shared code for lick direction. **Relates to Figure 4. A.** Schematic for aligning latent space. Spikes aligned to reward delivery from 5 mice underwent dimensionality reduction using PCA and subsequent alignment with multiset canonical correlation analysis (CCA) to identify shared subspaces across mice. **B.** Average variance explained by each canonical component across five mice. **C.** Total variance explained by 10 canonical components in free vs go trials. **D.** Temporal decoding generalization of rewarded lick direction from medullary spike data in remaining 4 mice. **E.** Individual confusion matrices for medullary cross animal decoding. Each confusion matrix represents the performance in held out mice. **F.** Performance of a model trained on 5 datasets with insertions in medulla pre-motor areas tested on a mouse with insertion in the hypoglossal motor nucleus. **G.** Cross animal decoding applied to aligned (PCA + CCA) cortical data demonstrates that shared code also exists in the cortex. **H.** Comparison of mean decoding accuracy in cortex and in medullary pre-motor areas. **I.** Absolute value of laterality canonical component weight for each principal component. Each dot represents the weight from one mouse. **J.** Weights of PCA for each unit.

### Inter-regional dynamics shift upon switch of motivational purpose

To examine the interactions between MOs and MY-mot during behavior, we analyzed data from sessions where we recorded simultaneously from both regions. Analysis of pairwise correlations between the firing rates of all units in the two regions revealed pairs with both strong positive and negative correlation coefficients (**Figure 5A),** across the entire recording session. Such correlations are consistent with the shared coding of different task epochs by individual units (see clusters in **Figure 2C–D**).

**Figure 5.**
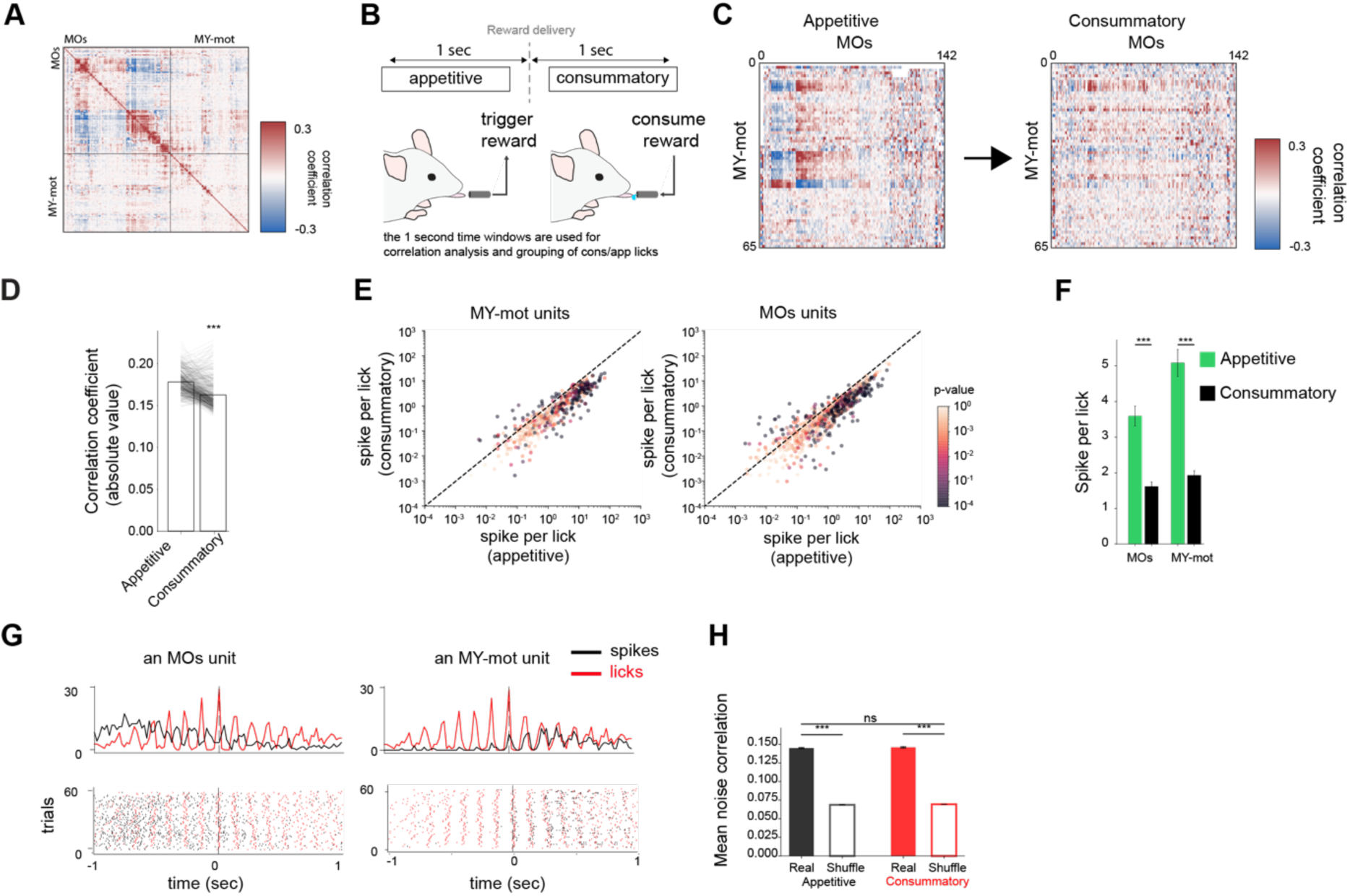
Pairwise correlations across regions during licking depend on motivational purpose. **A.** Pairwise correlation coefficients of firing rates between MY-mot and MOs units across “go” trials from an example session. **B.** Illustration of appetitive and consummatory licking, respectively before and after reward on “go” trials. **C.** Matrix of pairwise correlation coefficients between MY-mot and MOs firing rates in one-second time window before (appetitive) and after (consummatory) reward delivery. **D.** Absolute value of pairwise correlation coefficient of all unit pairs in appetitive and consummatory epochs. **E.** Scatter plot of spikes per lick of MY-mot and MOs units, color-coded with p-value (unpaired two-tailed t-test), during appetitive or consummatory phases. **F.** Spikes per lick of MY-mot and MOs units during appetitive (green) and consummatory (black) epochs. **G.** Example MOs and MY-mot units that have consistent bias towards either appetitive or consummatory licks. Only trials in which mice licked the same spout during both the operant response and subsequent reward consumption were included to control for lick direction. **H.** Pairwise noise correlation (NC) for appetitive (filled black) and consummatory (filled red) phases, overlaid with shuffle null distributions (unfilled).

To test how neural fluctuations between MOs and MY-mot units depended on behavioral context, we calculated pairwise neural correlations during appetitive and consummatory licking separately. We defined the appetitive and consummatory windows respectively as the 1 s immediately before and after reward delivery in “go” trials (**Figure 5B**). The appetitive window is defined as the 1 second before reward and therefore overlaps with the end of the odor period. We found that the strength of pairwise correlations between MOs and MY-mot diminished after reward delivery (**Figure 5 C-D, Figure S5**; absolute correlation coefficient: 0.225 ± 0.001 (n=326 epochs) pre-reward and 0.215 ± 0.001 (n=326 epochs) post-reward; p<0.001 Wilcoxon signed-rank test). This change also coincides with an overall decrease in the number of spikes per lick in the consummatory period (**Figure 5E-F**). This effect can be clearly observed even in the firing of individual units, demonstrating that motivational context can strongly affect unit activity in MOs–but also interestingly in the MY-mot (**Figure 5G**).

To test whether this pairwise correlation decrease reflects a change in baseline coupling or task-driven activation, we computed pairwise noise correlations (NC), which are absolute correlations computed after removing the mean response between all simultaneously recorded MOs and MY-mot unit pairs (n = 25,005 pairs across 3 datasets). Despite observed pairwise correlation decrease, NC did not change between the appetitive and consummatory phases (Figure 5H, Figure S5C, mean appetitive NC = 0.145 ± 0.004, mean consummatory NC = 0.146 ± 0.003; Wilcoxon p = 1.000 after Bonferroni correction; 50.0% of pairs had higher NC during consummatory phase). The change in NC across the transition was uncorrelated with the change in pairwise correlation at the pair level (Figure S5D, r = 0.002, n = 23,226 valid pairs), further suggesting these two measures are driven by separate mechanisms.

**Figure S5.**
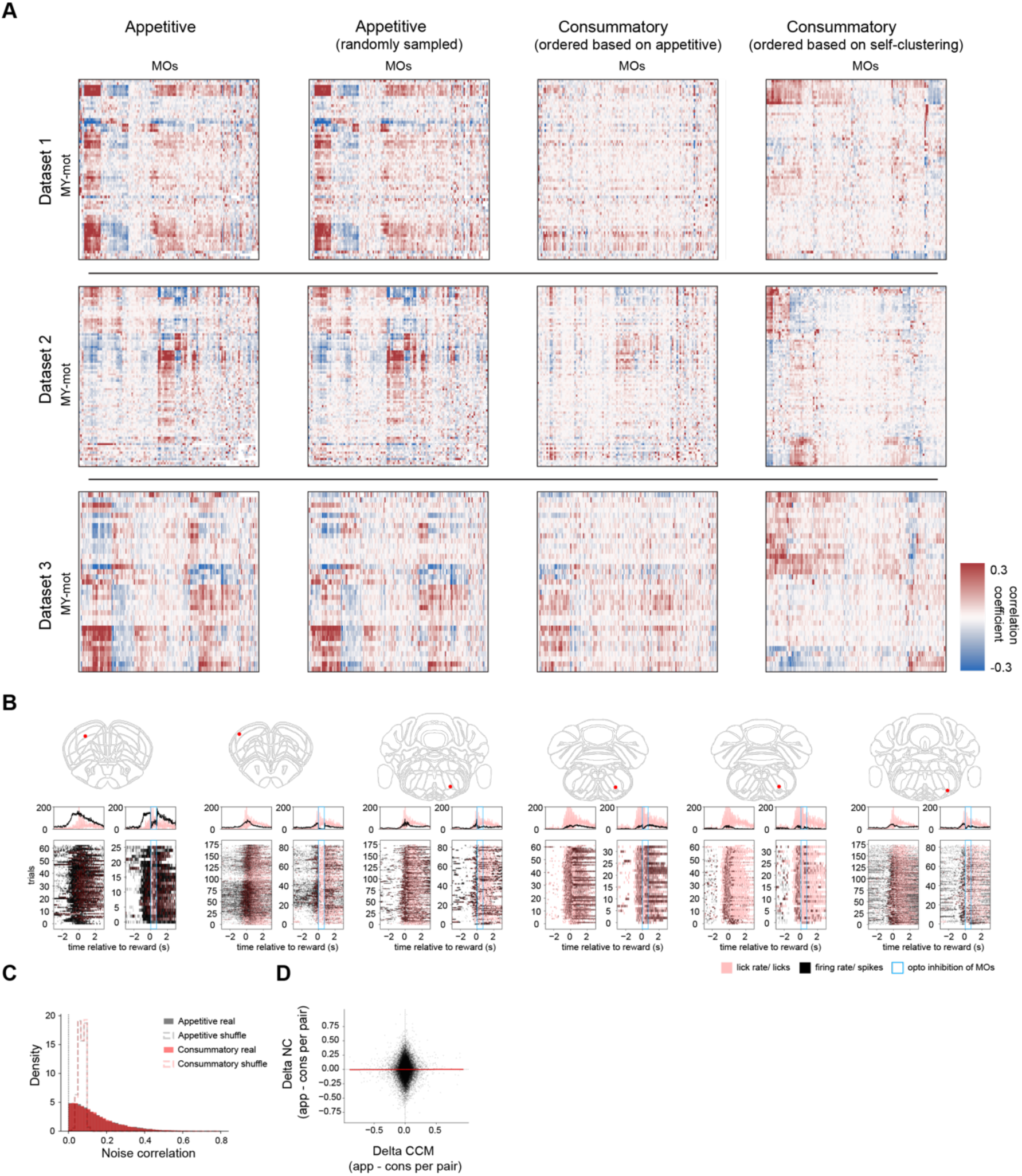
Encoding of purpose of licking between regions. **Relates to Figure 5. A.** Correlation coefficient matrix of MOs vs MY-mot. From left to right: sampling all appetitive epochs, sampling a random subset of appetitive epochs, sampling all consummatory epochs with ordering based on appetitive matrix, and sampling all consummatory epochs with ordering based on itself. **B.** Example MY-mot units modulated by MOs silencing. **C.** Distribution of pairwise NC (noise correlation) for appetitive (black) and consummatory (red) periods, overlaid with shuffle null distributions (dashed). **D.** Change in NC (ΔNC= NC_app_− NC_cons_) versus change in pairwise correlation (ΔCC= CC_app_−CC_cons_) for each MOs–MY-mot unit pair

### Cortico-brainstem communication subspaces depend on context

So far, we have shown that each of the recorded brain regions generates neural activity that is reflected by a few latent variables that are consistent across individuals (**Figure 4**). We have also found that simultaneously recorded pairs of neurons from MY-mot and MOs show common fluctuations (**Figure 5**), and that these correlations changed markedly during licking depending on the motivation of the animal, i.e., whether licking is appetitive or consummatory. These findings motivated us to investigate the properties of population activity in MY-mot and MOs during licking, and how they interact with each other depending on context. In particular, we assessed the properties of the “communication subspaces”^33^, which are defined as the subset of latent dimensions in one “source” region that can predict the neural activity of the other “target” region using a linear transformation.

To systematically assess these communication subspaces, we used cross-validated reduced-rank regression (**Figure 6A, Materials and Methods**). This analysis provides two complementary measures of communication: the dimensionality, namely the number of latent dimensions in the source region that correlate with the population activity of the target region, and the variance explained, i.e., how much variance in the target is accounted for by this communication channel. In order to compare the variance and dimensionality of inter-region communication subspaces, we also evaluated the communication subspace of a region with itself. To do so, we randomly assigned neurons of the same region to either a source group or a target group, and performed reduced-rank regression. Given that we included two areas in this analysis (MOs and MY-mot), we assessed four different communication subspaces (**Figure 6B**, for one example session): two internal or within-region subspaces (from MOs to MOs, from MY-mot to MY-mot), and two communication or across-region subspaces (from MOs to MY-mot, and from MY-mot to MOs).

**Figure 6.**
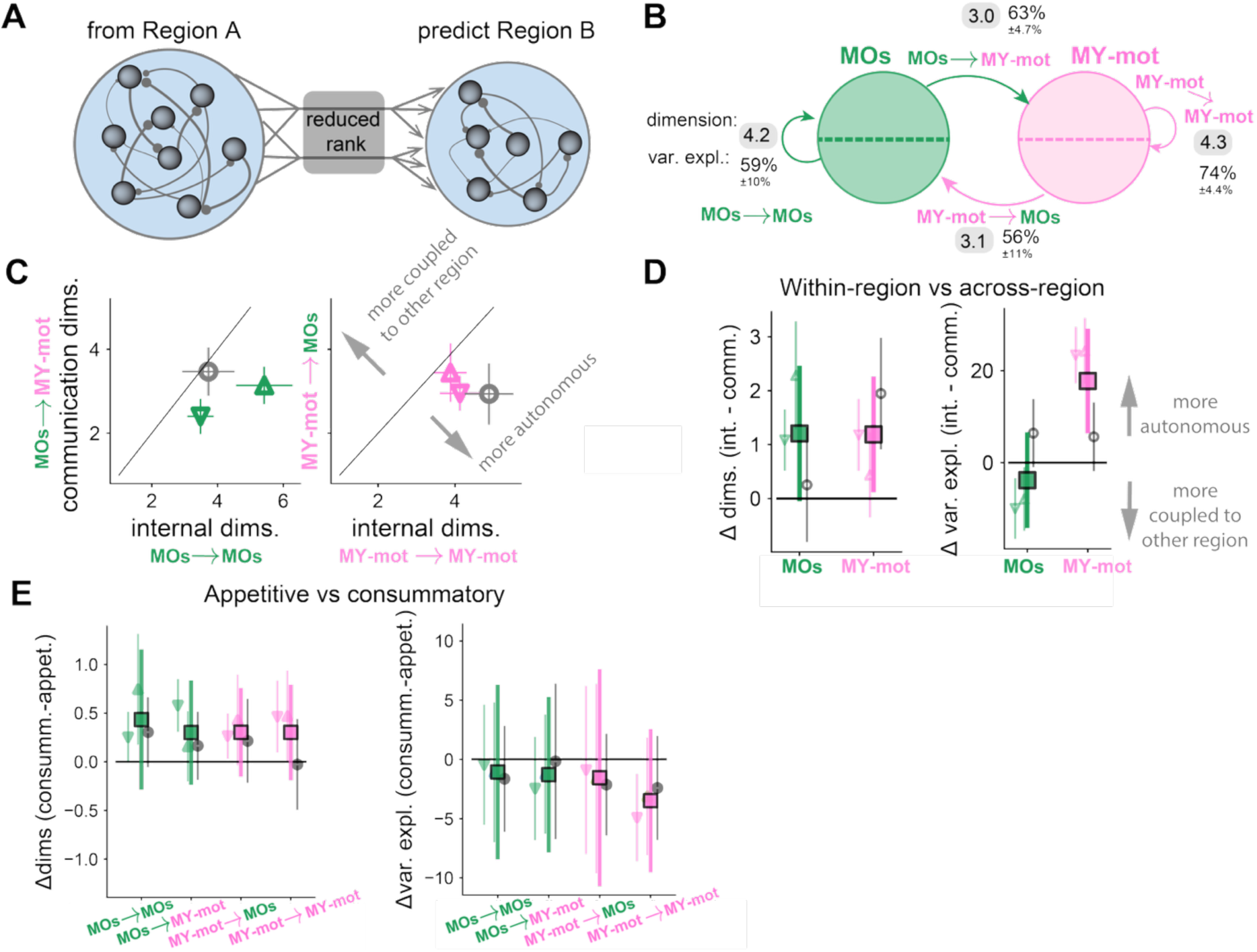
Communication subspace analysis reveals asymmetric interactions between cortex and the medulla. **A.** We applied cross-validated reduced-rank regression from a region A to linearly predict the activity in Region B. Regions A/B correspond to combinations of MOs and MY-mot. **B.** We estimated the average percentage of variance explained in the target area based on the source area, and the dimensionality of the communication subspaces. This example shows these measurements from one recording session. **C.** Dimensionality of communication subspaces in MOs and MY-mot. Each symbol corresponds to a different session. The gray circular symbol indicates the session where units in MY-mot lie primarily within the hypoglossal motor nucleus. The colored symbols indicate sessions where MY-mot units were primarily found in medullary premotor circuits, i.e., more anterior areas of the MY-mot containing premotor interneurons. **D.** Difference in dimensionality (left) and percentage of variance explained (right) between the within-region communication subspace and the cross-region subspace. The large square dots indicate the average across sessions. **E.** Difference in dimensionality and percentage of variance explained in each of the four communication subspaces during appetitive licking (before reward release in go trials) and during consummatory licking (after reward release in go trials). (ns: p >= 0.05; *: 0.05 > p >= 0.01; **: 0.01 > p >= 0.001; ***: p < 0.001)

During the time window around reward delivery, the dimensionality of the within-region subspaces was between 3 and 6 in MOs and MY-mot (**Figure 6C**). The cross-validated variance explained by these internal subspaces ranged between 50% and 80% (**Figure 6B, Figure S6**). Comparing these within-region characteristics with the cross-region communication subspace, we found that the dimensionality of the cross-region communication subspace was lower (**Figure 6D**). In particular, the cross-region communication subspace exhibited one fewer dimension than the within-region subspace. The differences in variance explained between the within- and across-region communication subspaces were particularly large in the medulla, especially in those sessions where the MY-mot neurons lay within the medullary premotor network. In those sessions, the latent variables within the medulla explained ∼75% of medullary activity but only ∼55% of cortical activity. These analyses revealed communication subspaces that show a clear asymmetry between the two regions. Specifically, MOs predicted MY-mot activity more accurately than vice versa.

Finally, given that the correlation structure changed with the trial epoch, we calculated and compared communication subspaces separately during appetitive licks and consummatory licks (before and after reward release, respectively). The dimensionality of both within-region and cross-region subspaces was higher during consummatory licks, but the total variance explained was reduced (**Figure 6E**). These results support the hypothesis that medullary and cortical networks produce more patterns of activity during consummatory licking and do so with a higher degree of autonomy.

**Figure S6.**
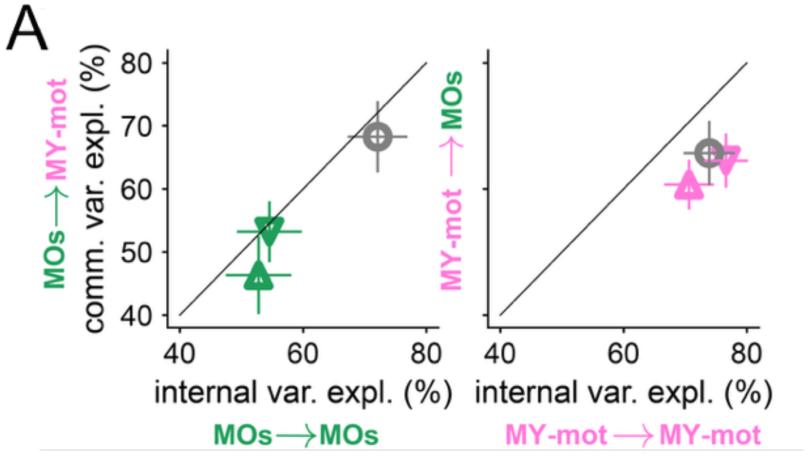
Variance explained by communication subspace analysis. **Relates to Figure 6. A.** Cross-validated variance explained in MOs and MY-mot, based on its within-area (internal) activity, or the across-area (communication subspace: comm.) activity. Each dot corresponds to a different session, the gray symbol corresponds to MY-mot units that lie primarily within the hypoglossal motor nucleus. Error bars indicate the standard deviation across cross-validation folds.

### Flexible interactions supported by multi-area recurrent networks

Communication subspaces are correlation-based measures of multiregional activity. To gain mechanistic insight into how units in MOs and MY-mot influence each other, we built recurrent neural network (RNN) models to reproduce the simultaneous neural recordings on a trial-by-trial basis (**Materials and Methods**). Such RNN models provide a mechanistic description of the data because, while reproducing the recordings, they also yield an estimated recurrent connectivity matrix and a set of input and output signals sufficient to generate the observed activity.

Our objective was to understand the recurrent interactions between cortex and medullary units during both “go” and “free” trials. But to successfully fit these data-driven mechanistic models, it was also necessary to provide a set of biologically-plausible external inputs to both the motor cortex and the medulla. Therefore, we chose a minimal set of external inputs that includes the licking and reward-related events we measured on each trial (**Figure 7A**). Due to this construction, most of the neural variability in the fitted RNN models was related to the recurrent interactions. We assumed two external input signals in each trial around the time of reward delivery: proprioceptive feedback to neurons in MY-mot, modeled here as the time-varying 3D tongue volume signal, and the reward signal, which is provided to both cortical and medullary units. To account for trial-to-trial variability, we further assumed that units may have a different initial internal state in each trial.

**Figure 7.**
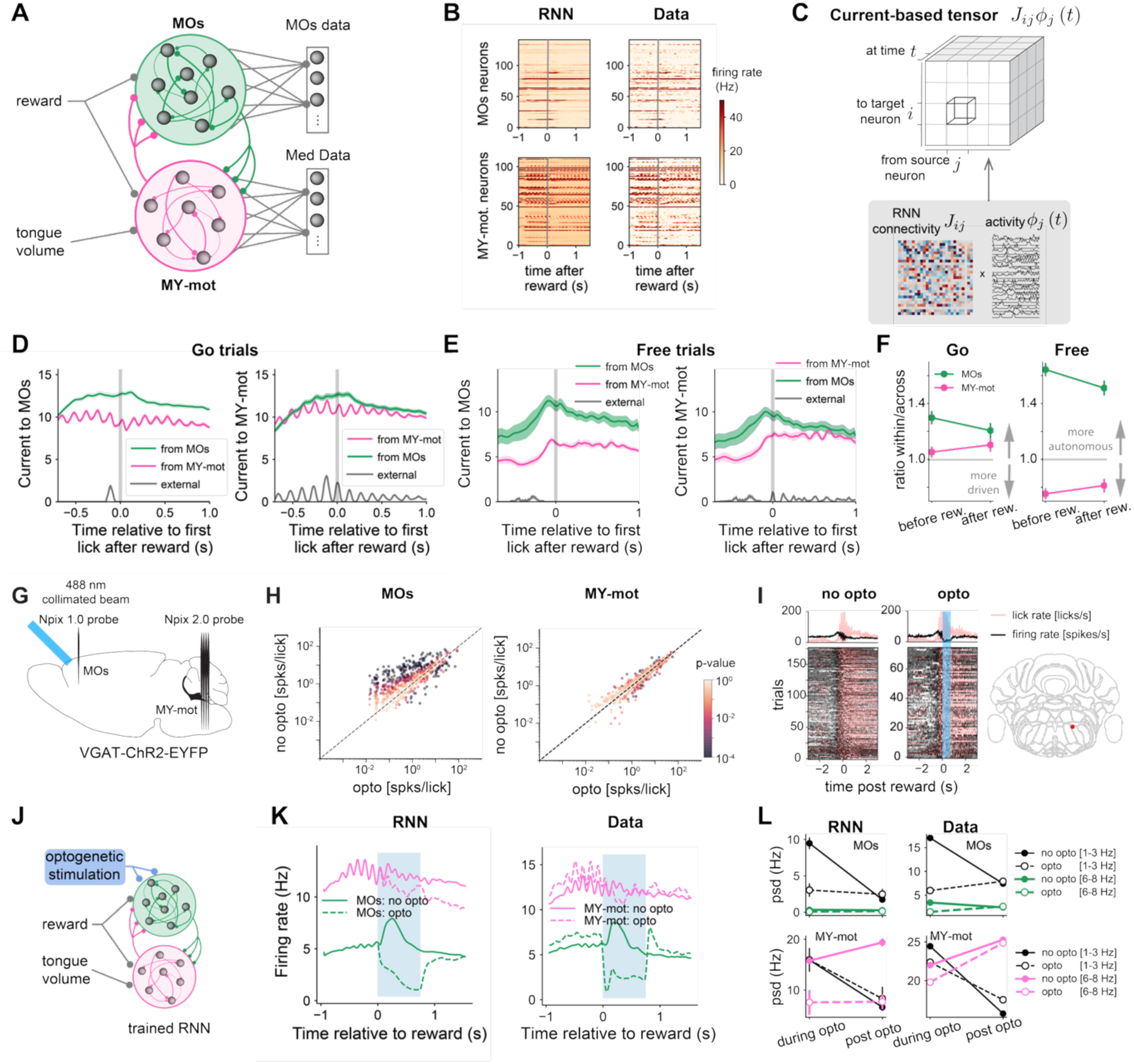
Current-based decomposition of activity around reward. **A**. Design of two-region recurrent neural network (RNN) trained directly on data. The reward signal is projected to units in both regions, while the tongue volume—a proxy for proprioceptive feedback or an afferent copy—projects only onto medullary units. Networks are trained such that linear combinations of the activity of MY-mot (medulla) units in the RNN matches the MOs (cortex) data. **B.** Example activity in the trained RNN during one trial (left), and neural recordings of that same trial (right). We focus on the activity preceding and following the reward delivery. **C.** Calculation of the current-based tensor from the RNN. The tensor encapsulates the directed interactions from every source neuron to every target neuron in the RNN, as a function of time. **D.** Average contribution to currents during go trials. We calculate the average current by taking the trial-average of the absolute value of currents from all source units in a given area to all target units in a specific area. Trials are aligned to the first lick after reward. In MOs, within-cortex currents are stronger than those coming from the medulla, with a slight decrease in magnitude following reward. In the MY-mot, both cortex and medulla-originated interactions are comparable in magnitude. **E.** Average contribution to currents during free trials. Currents ramp up right before lick initiation during free trials. Similar to go trials, currents within MOs are stronger than currents from the MY-mot projecting back to MOs. However, the current from MOs to MY-mot is stronger than the internal MY-mot currents. **F.** Ratio of current contributions: the within-region current divided by the across-region current contribution. **G.** Schematic of optogenetic silencing of MOs during dual recording from MOs and MY-mot. **H.** Spikes per lick of MOs and MY-mot units during opto stimulation vs without stimulation during the window of stimulation (0.7 second window after reward delivery). Color indicates p-values (unpaired two-tailed t test). **I.** Example MY-mot unit modulated by MOs silencing. **J.** Simulated optogenetic perturbation to the RNNs. An inhibitory stimulus is provided to neurons in MOs, reducing their firing rate. **K.** Average firing rate of units in MOs and MY-mot in the RNN and in neural recordings in trials with optogenetic stimulation. Blue indicates the period of stimulation. We only considered MOs units whose firing rate decreased during the stimulation period. **L.** Comparison of the power spectrum in low-frequency bands (black lines) and licking-related frequencies (colored lines), during the optogenetic stimulation and in the post-stimulation period. Comparison of RNN responses and data. Error bars show the average over 10 RNNs trained on the same recordings.

We then used gradient descent to fit the connectivity and single neuron parameters of the RNN (**Materials and Methods**, **Figure S7**). Through this process, our RNNs converged to a solution that largely reproduced the neural recordings, in particular its low frequency changes in firing rate (**Figure 7B**). In order to estimate the consistency of the RNN models, we trained 10 RNNs per recording session. To analyze the resulting RNNs, we applied a recently developed method, current-based decomposition (CURBD^34^). This decomposition can only be performed when both the recurrent connectivity and the single neuron activity are known and is therefore ideal for artificial neural networks. First, we calculated the tensor of time-varying currents that are sent from every source neuron to every target neuron in the network, by multiplying the synaptic weight between the source and the target with the source neuron’s activity (**Figure 7C**). In a second step, we averaged the contributions to the currents across units in the same region as the target neuron and across units from the other region.

The average contribution of currents to MOs and MY-mot varied over time and was modulated by reward (see **Figure 7D-F** for one example session, **Figure S7** for the additional sessions). During “go” trials, the largest contribution to the currents of cortical units stems from within-region recurrent connections. In the medulla, instead, the activity is driven by comparable contributions from cortical units and local interactions (**Figure 7D**). Similar current contributions are observed during consummatory licks in “free” trials (**Figure 7E**). When we compared the ratio between within-region and across-region currents, we found that, after reward release (during consummatory licks) the contribution of cortex to the medullary currents is reduced, indicating that the medulla is more autonomous during these intervals. In motor cortex, we also found a reduced influence of within-region currents during consummatory licks. These changes in currents during consummatory licks occur regardless of whether the animal is licking before reward, as they happen regardless of whether appetitive licks occur, as in “go” trials, or not, as for “free” trials (**Figure 7F**). Altogether, the CURBD analysis suggests that the motor cortex provides strong currents to the medulla, which are reduced after reward release, suggesting that medullary activity is more autonomous during consummatory licking.

We then analyzed the simultaneous neural recordings in MOs and MY-mot during optogenetic stimulation of inhibitory neurons in MOs (**Figure 7G**), to assess the predictive power of the trained RNNs to these optogenetic perturbations. As expected, inhibition of MOs following the reward release produced a large decrease in the firing rate of a large majority of recorded MOs units (**Figure 7H**). Interestingly, units in MY-mot experienced heterogeneous changes in their firing rate to MOs inactivation, showing only a small decrease in the average population firing rate (**Figure 7H**). The MY-mot responses were generally mixed, some units increased and others decreased their firing rate. Only a small subset of recorded MY-mot units were silenced by the optogenetic intervention **(Figure 7I**),

To then validate trained RNNs as mechanistic models of the interactions between MOs and MY-mot, we modeled an inhibitory stimulus onto the cortical units (**Figure 7J**) and simulated neural dynamics in response to this novel perturbation. By construction, the optogenetic-like stimulus in the RNNs reduced the firing rate in MOs, as in neural recordings (**Figure 7K**). We also observed an average decrease in firing rates in the medulla during the optogenetic intervention, in qualitative agreement with the recorded data. Additionally, in the post-stimulation period, the firing rates relax back to the unperturbed level in MOs and MY-mot. However, we also observed some differences between the data and the trained RNNs. For instance, the average firing rate takes longer to recover from the perturbation than in the neural data. This is expected in our simplified network models, which are built with rate-based neurons that react slowly to transient inputs^35^, and whose connectivity does not follow Dale’s law^36^.

To go beyond comparisons at the level of the average population firing rates, we compared the power spectral density of the RNN activity with the power spectral density of the neural data. We found that the changes in the power spectrum of single neuron activity during and after the optogenetic were qualitatively similar in the RNN and the neural recordings. We focused analysis on two power bands; for slowly changing firing rates (1-3 Hz), the signal power was reduced in MOs and MY-mot units in the post-optogenetic intervention period compared to the stimulation (**Figure 7L** and **Figure S7**), while for timescales corresponding to the licking rhythm (6-8 Hz), the power in the post-stimulus period remained constant or showed a small increase in MOs and MY-mot. The consistency of these results between RNNs and recorded data, in terms of changes in the average firing rate in the medulla and on their temporal statistics, validated use of RNNs to predict neural responses to novel perturbations.

**Figure S7.**
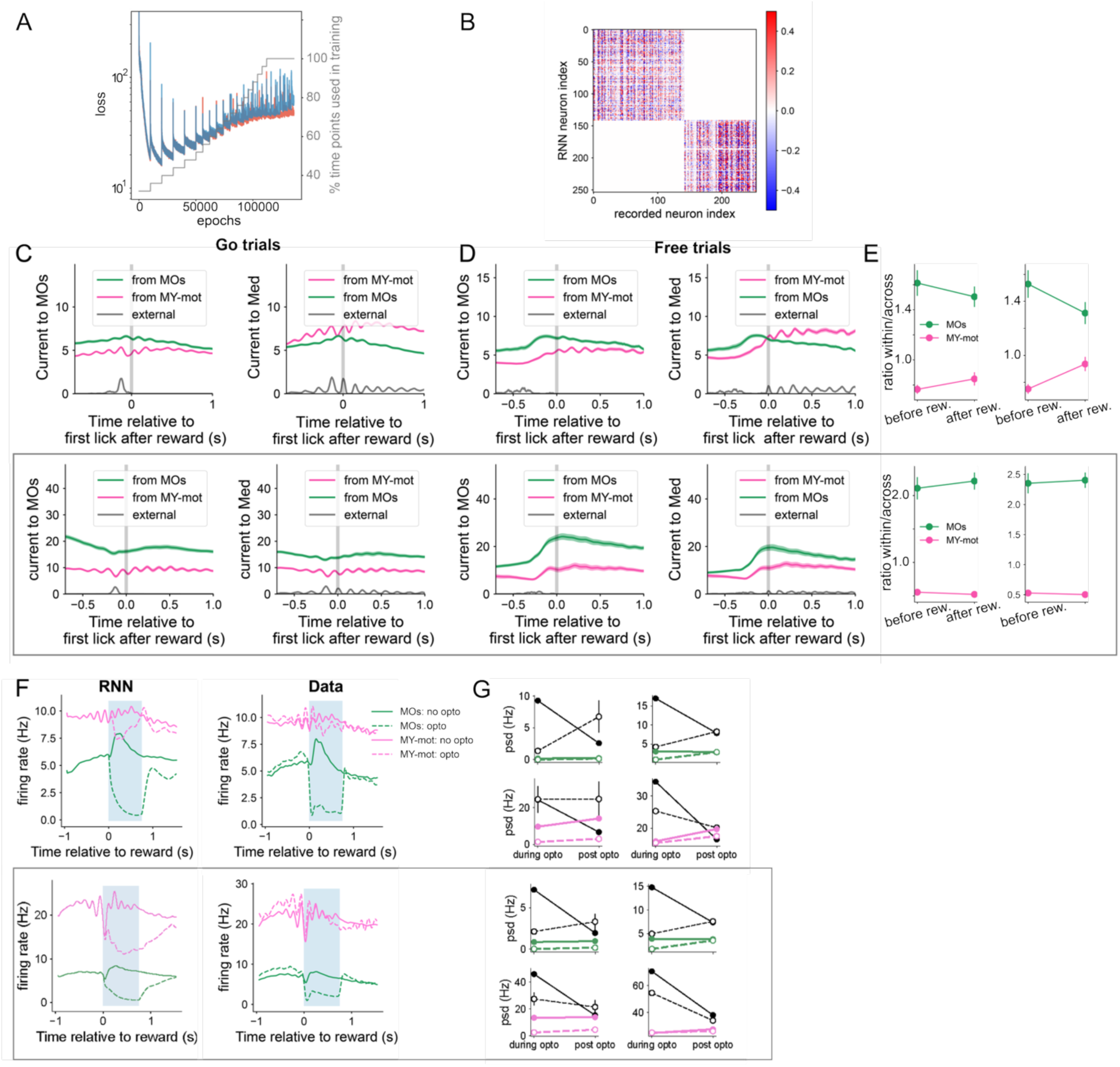
Trained RNNs and CURBD analyses across sessions. **Relates to Figure 7. A.** Loss function (blue and red) over the course of training for two different trained RNNs on the same session. In grey, the percentage of timepoints used in the training set. We gradually increased the number of timepoints used in training in order to obtain a better final performance. **B.** Linear readout matrix after training that maps the activity from RNN units to recorded single units. Each MOs (MY-mot) single neuron traces is a linear combination of the activity of the MOs (MY-mot) RNN units. **C-E**. Similar to Fig. 7 D-F, for a different session. Currents to MOs and MY-mot in “go” and “free” trials, as well as ratio between within-area current contribution divided by the across-area contribution. **F-G**. Similar to Fig. 7 G-H, for two additional sessions. The bottom row corresponds to the session where most MY-mot units lie within the hypoglossal motor nucleus. Average firing in MOs and MY-mot during optogenetic stimulation. Average changes in the power spectral density during optogenetic stimulation and post-optogenetic recovery.

## DISCUSSION

These experiments probed cortical influence onto the medulla during directed licking, and found that while cortex and medulla exhibit highly correlated activity patterns, cortical influence was stronger over the medulla than vice versa. In addition, the amount of cortical influence varied from moment-to-moment and depended on behavioral context–in particular, stronger when mice were engaged in operant, appetitive licking than in more automatic consummatory licking. Nevertheless, optogenetic inhibition of MOs during consummatory licking still had a significant effect on behavior–and on the firing of a subset of units in the MY-mot.

### Interactions between MOs and MY-mot change with the motivational context of movement

Over the multiday timescales associated with mice learning to perform a motor task, the causal relationship between cortical activity and movement is known to weaken^6,8,37^. Over shorter timescales, cortical influence over movement is also known to be variable. For example, it is thought to be particularly instructive during the activation of certain limb muscles ^38^ and in making corrective submovements of the tongue that are needed to reach a waterspout ^5^. These findings are consistent with the hypothesis that cortical engagement is strongest in the context of a need to rapidly adjust behavior due to uncertainty in the environment ^6^. Our results are generally supportive of this idea. We recorded from highly-trained mice that had learned to perform a behavioral task wherein they could make any sequence of directional licks in response to a cue in order to retrieve a reward. Over the course of learning, each animal developed a highly stereotyped licking pattern in response to the “go” cue. But as the reward was dispensed from a random spout on each trial, mice had to redirect their licks in new directions that were impossible to predict. Therefore, in our task, the time surrounding the reward delivery reflected a period of maximum uncertainty, requiring readiness to modulate behavior. We identified this pre-reward appetitive period as a period of strong cortico-medullary coupling.

Our findings go beyond previous work that focused almost exclusively on the direct relationship between cortex and behavior during these epochs of high cortical engagement, rather than on the relationship between cortex and the brainstem circuits that drive motor nuclei ^4–6,15,38^. By leveraging our simultaneous recordings from MOs and MY-mot, we were able to assess the interaction strength between these areas across two motivational contexts. Across multiple analyses, including pairwise correlations, reduced rank regression, and as modeled using RNNs, we observed stronger cortical influence during the appetitive epoch than during the consummatory epoch. This was true even though the licking actions during the appetitive period were highly consistent from trial to trial within a session, while consummatory licks strongly differed in direction from trial to trial. On the other hand, we found noise correlations to be stable during the consummatory-appetitive transition, suggesting that the decrease in coupling between two regions is driven by changes in activity that remains constant across trials, rather than trial-to-trial variability. We note that we did not set out to analyze “communication through coherence” in this work^39^, instead focusing exclusively on spiking and firing rate correlations. But future work could examine whether cortico-medullary coherence in either the gamma band, or in the frequency bands associated with licking or breathing^40^, might also play a role in controlling the amount of influence that cortex has over movement during various motivational contexts.

Of note, we refer to licks made post-reward as “consummatory”. But we do not mean to imply that the presence of reward itself changes the neural control of the tongue. Indeed, ventral tegmental area (VTA) stimulation experiments during licking^6^ showed that the mere presence of a reward-related signal is not sufficient to engage the cortex. Instead, by “consummatory” we simply mean that the licking behavior leads directly to reward: the mouse does not need to consider any stimuli beyond the reward itself once it has felt the water droplet on its face and has begun to consume it. Future work could explore the limits of this distinction in a greater diversity of motivational contexts, including in settings wherein tradeoffs exist between reward acquisition and an aversive outcome. For example, it has been shown that direct optogenetic pacing of elevated heart rate imposes an anxiety-like behavioral state, such that even thirsty mice will substantially delay licking for water reward if a low (10%) risk of mild shock exists—but a similar risk of shock does not delay licking in the non-elevated heart rate condition^41^. The medullary cardiovascular center contains distinct regions involved in sensing and controlling heart rate, including the nucleus of the solitary tract, which communicates afferent cardiac information to insular cortex in the forebrain as well as to oromotor nuclei in the medulla. It will be interesting to explore whether aspects of the neural control of the tongue are altered in this or other distinct contexts of altered motivational state.

### Population-level representations for specific motor actions in the medulla

In sum, we have shown that cortico-medullary interactions change on short timescales that are motivationally-salient to the mouse. Despite this time-varying cortical influence, the medulla must implement a structured motor program to convert descending input into action. Conversion of constant descending command inputs into rhythmic motor outputs is accomplished by central pattern generators (CPGs)^42^. Such circuits exist in the brainstem, where they are thought to coordinate a wide range of orofacial behaviors including breathing, licking, swallowing, suckling, chewing, and vocalization, often by providing monosynaptic input to motor neurons^17,20,22,42^. Recent work in turtles suggests that spinal CPGs that control rhythmic limb movements employ low-dimensional rotational population dynamics where spinal cord interneurons participate in periodic patterns of population activity at specific phases^24^. Consistent with these results, large-scale recordings from the medulla of mice during licking have revealed that many medullary interneurons are also phase locked to the lick cycle^3,13,43^.

A central unresolved question, however, is how multiple distinct motor patterns can be generated by the same oscillator population. Here we examined whether the timing of phasic bursts, independent of spike frequency, is itself sufficient to represent distinct movements. First, we showed that in the MY-mot, neuronal phase offsets over the lick cycle depend on lick direction. Second, we showed that lick direction can be decoded directly from the distribution of phases across the population, indicating that the organization of rotational dynamics itself encodes task variables. Together, these results impose new constraints on the mechanisms by which CPG circuits can generate subtly distinct motor outputs.

The presence of strong direction-specific phase codes across animals further suggests that there may exist a population-level code in the medulla that represents licking movements with distinct kinematics in an invariant manner across animals. This would be consistent with a study of primate motor cortex showing that limb movements could be reliably decoded across distinct neuronal populations taken from different individuals by aligning neuronal population dynamics using CCA^31^. Here we extend these findings to the mouse medulla to show that licking kinematics may also be generated by broadly similar neuronal population dynamics across different anatomical locations in the medulla–and even across mice. These results suggest that the medullary premotor regions may implement a structured, low-dimensional plan for specific actions, one that is conserved across animals. The failure of this shared code to generalize to hypoglossal motor neurons further emphasizes that this shared latent space is more than just a simple surrogate for muscle activity or biomechanics.

### Analysis of multi-region interactions

Our analyses of simultaneously recorded neuronal activity data from cortex (MOs) and from a postsynaptic region in the medulla (MY-mot) revealed that medullary activity was strongly modulated by MOs activity while mice were engaged in the task, particularly around the time of reward release, and in a manner that depended on the animal’s motivational state. Yet, the causal direction of these interactions is difficult to establish: MOs activity also co-varied with activity changes in MY-mot. Such bidirectional correlations may reflect direct anatomical connections, for example, motor fibers from MOs directly project to the MY-mot^13,44^ and the medulla provides feedback to cortex via the thalamus^45^. At the same time, shared fluctuations could arise from shared inputs, such as from unrecorded brain regions or sensory signals such as proprioceptive feedback.

As an initial step toward characterizing shared activity across regions, we used a communication-subspace method^33^. This cross-validated approach provides two complementary measures: how much variance in a target region can be explained by a source region’s activity, and how many latent dimensions account for that variance. Jointly, these two statistics may provide hints about the directionality of the interactions. For instance, MY-mot activity was better predicted by MOs than MOs predicted by MY-mot, suggesting a stronger influence of top-down MOs input onto MY-mot than from bottom-up feedback pathways. Nevertheless, a full exploration of inter-area interactions must go beyond identifying shared latent variables, and instead provide mechanistic hypotheses about how information is transferred and transformed across regions^46,47^.

To achieve this outcome in a manner beyond simple correlational analyses, and to study the nonlinear recurrent dynamics underlying these interactions, we trained multi-region RNNs that reproduced neural recordings using the current-based decomposition framework (CURBD^34^). This approach allowed us to decompose each model neuron’s activity into weighted contributions from all presynaptic units. This is possible because the full connectivity of the model is known. While these models are powerful, their reliability remains debated, and depends critically on the recorded data and additional modeling constraints^48–50^. In practice, we view these approaches as hypothesis-generating tools that can be experimentally tested with novel data. Accordingly, we compared the RNN responses to inhibitory stimuli in the motor cortex, similar to our optogenetic cortical inhibition experiment. Although some differences were observed between the model and the data, the RNNs reproduced key features, notably at timescales associated with licking (∼8 Hz) and behavioral context (∼1 Hz). This level of experimental validation provides an estimate of the degree to which RNN models can be relied upon, and suggests that these models may indeed be useful for guiding future experimental protocols.

Together, our multi-region neural recording experiments, combined with optogenetic perturbations and multi-region RNN models, revealed new insight into the context-dependent contributions of cortex to orofacial behavior. They also set the stage for future studies that further dissect the specific contributions of various descending command pathways across the central nervous system in the generation of movement.

## MATERIALS AND METHODS

### Experimental Model and Subject Details

All animal procedures were conducted in accordance with protocols approved by the Stanford University Institutional Animal Care and Use Committee (IACUC) and guidelines of the National Institutes of Health. Both male and female mice were used, aged 2 - 8 months at time of first recording (typically about a month after beginning training, see “Animal Behavioral Training” below). Mice were group housed in plastic cages with disposable bedding on a standard light cycle until headbar implantation, at which time they were moved to a 12 hr reversed light cycle room. Following recovery after surgery, mice were put on a water restriction schedule where they received approximately 1 mL/day of water. All experiments were performed during the dark period. All experiments were performed with mice from the VGAT-ChR2-EYFP strain (JAX stock #014548) bred on a C57BL6/J background.

### Animal behavior apparatus

As described previously^51^, a microcontroller-based real-time behavioral system (Sanworks, Bpod State Machine r1) was used to control delivery of stimuli and water reward. Three independent waterspouts (Dispense All brand blunt dispensing tips, 22 gauge, 2” length) were manually bent and connected via luer locks to a custom manifold (fabricated by ProtoLabs, CAD model available upon request). This resulted in the middle spout being centered on the mouse’s incisors with its tip positioned at a distance of approximately 3.2 mm, and the two side spouts being offset at an angle of approximately 60 degrees with the left spout at about 2.9 mm and the right spout at approximately 2.7 mm. The precise position of the spout manifold relative to the mouse’s incisors was reproducibly achieved on each experimental session as it was mounted on a post conveyed by motorized translation stages (Thorlabs MTS25/M-Z8) and aligned to a region of interest (ROI) in the bottom up camera view (see details later in this section). The mouse was affixed to a custom head fixation device identical to previously described^51^ but with a modified headbar design (fabricated with stainless steel by Protolabs, CAD model available upon request) that exposed the posterior skull over the cerebellum (see Surgical Preparation).

Water was delivered via a gravity-fed reservoir attached to the lickometer and controlled by a quiet solenoid valve (Lee Valve LHDA1231115H). For olfactory stimuli, an olfactometer was constructed using pure odorants (ethyl acetate and 2-pentanone, Sigma) diluted to 4% v/v in paraffin oil (Sigma) and pressurized with air (1 L/min). Two 3-way valves (NResearch #161T031) controlled odor delivery, with the normally open port connected to a blank vial and the normally closed port connected to an odor vial. Odor delivery was controlled by actuating solenoid valves to switch airflow away from a blank valve and to the odor valve. Odors were delivered through a Teflon tube (NResearch, #TBGM109 with #102P109-10 connectors), placed approximately 1 cm from the mouse’s nose. Two color computer vision cameras (Basler acA1300-200uc), each fitted with a 25mm fixed-focal length lens (Edmund Optics #59-871), were used to measure the animal’s face from two orthogonal angles: from the right side, and from the bottom. The surface of each camera lens was positioned about 27 cm from the mouse’s head. Video acquisition was performed using custom software at 200 Hz (Python, using the pypylon library). A small red LED was placed in front of each camera, and was synchronized by Bpod to flash for 0.5 seconds, 2 seconds before odor onset (on “go” or “no go” trials) or reward (on Free trials). Valve openings (for odors and rewards), LED flashes, and optogenetic pulses were all controlled programmatically with the Bpod.

Licks towards each spout were optically detected in real time. To achieve this, we wrote custom Python software that processed ROIs positioned over each spout tip in the bottom up camera view of the mouse’s face. Our lick detector relied on a threshold that detected the unique color of the tongue relative to the color of each spout within each ROI. Detected licks were relayed from the computer via an Arduino Uno that sent event times to the Bpod. On each trial, and in each experimental session, we measured the round trip latency for a Bpod-triggered red LED flash to be detected by the Python software and relayed back to the Bpod. This typically took less than 0.02 s, or 4 camera frames at 200 Hz.

All timing signals were acquired at 20 kHz using a National Instruments PXIe-8381 card to measure frame onset signals from the behavior cameras, the trial start signal from the Bpod, and the synchronization signal from the IMEC Neuropixels PXIe acquisition module.

### Animal behavioral training

After waiting at least a week after surgery, mice were rationed to 1 mL/day of total water while ensuring that they maintained > 80% of their pre-deprivation weight. Mice were then handled for several days and habituated to head fixation. Following this, they began a four-stage behavioral training process that took between two and three weeks from the first day of training until behavioral performance had stabilized and the animals were ready for recording.

In the first stage, mice were head fixed and placed in front of our three-spout apparatus. On each trial, they were presented with an odor (ethyl acetate) for 1 s. A water reward of approximately 1 µL was automatically dispensed 500 ms after the odor turned off. Once mice began to reliably lick in response to the odor cue, the mice moved to the second stage of behavioral training where the mice had to lick to a spout during the 500 ms time window following the offset of the odor in order to trigger reward. We initiated the third stage of training once mice reliably triggered rewards on these “go” trials. On these sessions, we additionally introduced two new trial types: “no go” trials (20% of all trials), where a second odor (2-pentanone) was delivered and no reward was delivered regardless of the animal’s behavior and “free trials” (20% of all trials) where reward was randomly delivered from one of the three spouts with the same timing as an odor would be normally delivered on a “go” trial. The final stage of training, which represents the full task, was identical to the previous stage but with the addition of optogenetic stimulation (see transcranial optogenetics, below). All neural recordings were conducted during this final variant of the behavioral task after animals had achieved stable performance on the task. This was defined as having two experimental sessions with average performance > 80% between trials 50-100 on the full task. Performance on “go” trials late in each session declined as the mice became sated.

To summarize the timing of the stimuli on the final stage of the task, on “go” trials, the olfactory stimulus was on for 1 s. Then there was a delay period for 0.5 s. Following this delay, the mouse needed to lick to any spout to trigger a reward from a random spout. If the mouse does not lick during this post-delay window (of up to 1 s), then they did not receive a reward. “No go” trials had the exact same timing pattern, but no reward was dispensed regardless of the mouse’s actions. “go” and “free” trials were defined as correct if the mouse triggered and consumed a reward within 4 seconds of it being dispensed. “No go” trials were defined as correct if the mouse did not lick within 4 seconds of the time in which a reward would have been released on a “go” trial (**Figure 1B)**.

### Surgical Preparation

Before behavioral training, all mice (all VGAT-ChR2 animals as described above) were surgically implanted with a headbar following a procedure similar to previously described^52,53^. Prior to surgery, mice were placed into a stereotaxic frame and were anesthetized with isoflurane (1-2%). Their eyes were coated with ointment (Puralube), and then Buprenorphine ER (0.5 mg/kg) was administered. After removing fur and cleaning the scalp with Betadine and isopropanol, a midline incision was made along the dorsal surface of the head periosteum was removed, and the skull was cleaned, dried, and leveled in the stereotaxic frame. Then a custom stainless steel headbar (shaped like a circle over the dorsal surface of cortex; fabricated by Protolabs) was cemented to the skull using clear dental cement (Parkell Metabond). We then marked out the following coordinates on the skull surface with a surgical pen for transcranial optogenetic stimulation and/or neural recording: 2.5 mm anterior, 1.5 mm lateral to bregma for anterolateral motor cortex (ALM/MOs) and approximately 5.8 mm posterior and at 1 mm lateral to bregma to primarily target the intermediate reticular nucleus and other locations in the motor medulla (MY-mot). After marking, an additional layer of clear dental cement was then applied to cover the dorsal surface of the skull in order to better connect the circle-shaped portion of the head bar to the skull surface–and to reinforce the skull’s integrity over time.

On the week of the first Neuropixels recording session, mice were again mounted on the stereotax under isoflurane anesthesia. A craniotomy was then drilled over the cortex (typically at approximately 1 mm anterior and 3.5 mm posterior to bregma). Next, the stereotax was used to place a gold grounding pin affixed to a platinum-iridium wire (0.002 mm diameter, A-M systems) was surgically implanted into the brain and cemented down with dental cement.

On each neural recording day, mice were placed on the stereotaxic frame under anesthesia, and a new craniotomy (or set of craniotomies) was drilled. Mice were moved to the head fixation system on the recording rig immediately after recovery from anesthesia.

### In vivo electrophysiology

We used single-shank Neuropixels 1.0 probes^54^ to record from locations in the cerebral cortex and quad-shank Neuropixels 2.0 probes^55^ to record from targets in the medulla. On different sessions, recordings were made from each hemisphere. Acquisition was performed using SpikeGLX. Data were sampled at 30 kHz and synchronized to timing signals acquired at 20 kHz from our National Instruments PXIe-8381 card. Craniotomies were drilled on the day of each recording and were kept moist by frequent application of phosphate-buffered saline (PBS). Craniotomies made to facilitate insertion of quad-shank Neuropixels 2.0 probes were at least 1 mm in diameter. Those made for single-shank probes were significantly narrower. Craniotomies were typically centered near the skull markings that were made during the earlier surgery used to attach the headbar, with some intentional variance over the medulla in order to target additional locations along the anterior/posterior axis. Prior to probe insertion, each probe was dipped into a solution of CM-DiI Dye (ThermoFisher Invitrogen #C7000) in ethanol to label insertion tracks. Probes were inserted vertically using two Sensapex uMp-3 micromanipulators at approximately 3 µm/s, as slow insertion rate is known to improve unit yield^56^. Insertion depth was monitored in real time by zeroing the probe location at the surface of the brain and then monitoring the local field potential (LFP) and real-time spiking data during insertion. After each experiment, probe insertion coordinates were determined histologically (see below). For probes inserted into hindbrain, gross differences in spiking activity could be observed between the cerebellum, the fourth ventricle, and the medulla. Following each recording, craniotomies were closed with Kwik-Cast (World Precision Instruments). Probes were then cleaned with Tergazyme solution and were reused.

### Transcranial optogenetics

A single behavioral apparatus was used for all behavioral training and neural recording. This system was equipped with a 450 nm laser diode light source (Doric LDFLS-450/075, 75 mW). The laser was coupled through an optical fiber system and a splitter onto two pairs of mirrors, collimators, and irises (Thorlabs) in order to generate two ∼4 mm-diameter spots of light on the skull, positioned approximately 33 cm below each collimator. Power at the skull surface was approximately 18 mW per spot (≈1.43 mW/mm²). Before each behavioral session, VGAT-ChR2 mice were head-fixed and each spot was manually aligned to MOs using a target marked onto the skull (see “Surgical Preparation”). All stimulation was performed bilaterally, targeting the same cortical area in each hemisphere during each experimental session.

Optical stimulation was performed on a randomly-chosen 30% of all trials during each session. On stimulation trials, the laser was continuously activated by a square pulse to its maximum power, at the onset of reward, for 750 ms. In order to control for potential any behavioral effects that have resulted from the mouse observing the optogenetic light source, on all trials (both stimulation trials and non-stimulation trials), a blue masking LED placed in front of the mouse’s face was also activated for 750 ms at the onset of reward.

### Brain clearing, electrode track reconstruction, and atlas registration

Following each experiment, mice were euthanized and were first transcardially perfused with ice-cold phosphate buffered saline (ThermoFisher), and then with a 4% paraformaldehyde solution (Electron Microscopy Services). Brains were dissected out of the skull and were then subjected to a previously-described chemical clearing procedure^53^. Cleared brains were then transferred to index-matching solution (ethyl cinnamate, ThermoFisher) and were imaged to acquire data with 2 x 1.794 x 1.794 µm-sized voxels using a LifeCanvas SmartSPIM light sheet microscope. Images were acquired in two color channels: a 488 nm excitation channel (for atlas alignment), and a 561 nm excitation channel (where the CM-DiI tracks left by each probe insertion were visible).

These imaging data were then spatially-downsampled to 25 µm isotropic voxels. The autofluorescence channel was then computationally registered to the Allen Brain Atlas reference data (CCFv3)^29^ using the BrainReg python package^57^. Once our raw data was warped into the common reference atlas space, we used the BrainReg napari plugin to trace each electrode track visible in the signal channel. We then used custom python scripts to align the Neuropixels electrode map to each traced track (following a similar procedure for both Neuropixels 1.0 and 2.0 probes). To do this we superimposed the Neuropixels electrode map onto each track and then used a custom GUI to manually identify a scale factor that rescaled the Neuropixels electrode maps appropriately for each animal. Using our GUI, we identified an electrode map scale factor for each mouse such that features in the average local field potential (LFP) power data from each recording were aligned with atlas-defined boundaries between brain areas, ventricles, and surface of the brain. Once the electrode maps were aligned to each track, we then assigned each electrode to a brain area in the CCFv3. The location of each unit was defined to be the unit where its spikes were most prominently detected by Kilosort (see ‘Spike Sorting’ below). All analyses focused on units that were defined to be in either secondary motor cortex (‘MOs’ in the CCFv3 brain area taxonomy), or motor-related areas in the medulla (‘MY-mot’).

## Quantification and statistical analysis

All analyses were performed using custom Python scripts that will be made available upon publication.

### Behavioral video analysis and tongue tip tracking

Online detection of licking was accomplished as described above, but all subsequent offline analysis was performed instead using a custom 3D tongue tracking pipeline. We developed a pipeline roughly consistent with previous work^5^. First, we synchronized our two 200 Hz color Basler cameras using frame exposure time information acquired using our DAQ card in conjunction with red LED pulses that occurred synchronously at the onset of each trial. These timing signals were also used to synchronize Bpod behavior events with movement signals extracted from the camera data. Our two cameras were positioned orthogonally relative to one another: one looking upwards at the mouse’s head from below, and another positioned to the right side of the animal’s head. Before further processing, both videos were each downsampled and cropped to images of 256 x 256 pixels. Next, we interactively defined circular regions of interest around the three lick spouts and a tongue “center” position (centered in the middle of the animal’s mouth) for each dataset. Defining these common locations between the two camera views allowed us to spatially align the two videos to each other.

We then performed tongue segmentation independently on the side and bottom view using UNets. Both models used a similar architecture (RGB input, 2 output channels, padding = “same”, base filters = 16; model based on https://github.com/jakeret/unet) with depth = 4 for the bottom-view model and depth = 5 for the side-view model. The bottom model was trained on a corpus of 1800 total frames where 940 frames had visible hand-labeled tongues and the rest had no tongue visible. The side view was trained on 1200 total frames where 320 had tongue images visible. These frames were taken from videos taken from three different mice. Simple data augmentation was performed by flipping each frame to effectively double the amount of training data. None of these training datasets were used in any subsequent analyses. Models were trained for 30 epochs and we used a 70% train, 15% test, 15% validation set split. Final model weights were saved and were later loaded up by our analysis pipeline and applied to all experimental datasets.

Using videos segmented with these UNet models, 2D tongue statistics like tongue centroid and tongue area were computed independently using each view. We also computed 3D statistics by aligning the results of tongue segmentation between the two cameras to reconstruct a 3D convex hull that encompassed the tongue on each frame. These convex hulls defined the tongue volume, and by following a previously published procedure^5^, it also identified the tongue tip on each frame. No camera calibration (e.g. calculation or correction of camera intrinsics/extrinsics) was performed beyond the alignment of the views. For visualization, 3D tip trajectories were Kalman-smoothed (pykalman; EM 5 iterations).

We further cleaned the detected lick events by setting frames in the 1D “tongue volume” trace with no visible tongue to zero and smoothing it with a Gaussian kernel (σ = 1 frame). Candidate licks were then defined as local maxima in the tongue volume that were separated by ≥10 frames at 200 Hz (≥50 ms; ≤20 Hz). Spout assignment was performed from the bottom view; if an event overlapped both the middle and a lateral spout, it was assigned to the middle spout (the tongue commonly occluded a lateral spout en route to the farther middle spout). We required all retained spout contact events to have agreement from both camera views, retaining only events with spout contact in both side and bottom views within a ±10 frame (50 ms) window; single-camera contacts were placed in an “unassigned” bin. Analyses therefore used four categories (left, middle, right, unassigned).

#### Spike sorting

Our raw electrophysiological data were processed using a custom pipeline based on Jennifer Colonell’s ecephys_spike_sorting package (Janelia Research Campus). Data was preprocessed using utilities included with SpikeGLX (Bill Karsh, Janelia Research Campus). Briefly, data was filtered and preprocessed using CatGT and C_Waves. The preprocessing pipeline parameters were identical between mice except CatGT ran with global common average referencing (’gblcar’) and artifact removal for 30/37 recordings and using local common average referencing (’loccar’) for 7/37 recordings. Data was then synchronized with behavior events (TPrime) and spike sorted using Kilosort3^58^ (https://github.com/MouseLand/Kilosort/; git commit hash e1b4d6114d2bed8b5e01f56785a8314d8882bc2c). Spike sorting quality metrics^59^ were then computed with the module included with ecephys_spike_sorting. Owing to differences in the refractory period between neurons in the medulla and cortex, the ‘isi_threshold’ parameter (which is the time window used for ISI violations) was set to 1.5 ms for medulla recordings and 2 ms for cortical recordings. We retained only units that had a low rate of interspike interval violations (‘isi_viol’ < 0.1), high signal-to-noise (‘snr’ > 1.5), and at least a moderate number of spikes (‘nspikes’ > 500). These remaining units were then manually curated using Phy (Cyrille Rossant, International Brain Laboratory). The resultant list of putative single unit “neurons” was used in all subsequent analyses.

### Intended lick spout

For licks that could not be assigned to a single spout from both camera views (”unassigned” licks), the intended spout was inferred from the 3D tongue volume. For each lick, we computed the minimal Euclidean distance between any voxel of the tongue volume and each of the three spout positions across the frames surrounding the lick event (±15 frames, ±75 ms), and assigned the spout with the smallest distance as the intended spout. Licks for which two or more spouts shared the same minimal distance, typically because the reconstructed tongue volume reached more than one spout, were considered ambiguous and excluded from analyses of lick direction (**Figure 1M–N**; 6.6% of licks in laser-off and 15.5% in laser-on trials, 0–0.7 s after reward).

### Half-peak width

In **Figure 2G**, for each peristimulus time histogram (PSTH), the peak response was identified. The signal was then traced leftward and rightward from the peak until it fell below 50% of the peak amplitude. The time interval between these two points was defined as the half-peak width and was measured in milliseconds.

### Response ratio

To determine the response ratio of each neuron, each lick was assigned a time window spanning from the midpoint between that lick and the preceding lick to the midpoint between that lick and the following lick. For licks whose preceding or following lick occurred more than 0.29 s away (average lick interval within lick bouts), a fixed half-window of 0.145 s was assigned instead. The response ratio was then calculated as the fraction of lick-associated time windows in which the unit fired at least one spike.

### Linear regression of tongue kinematics

Datasets containing more than 30 units in each of both regions were included in linear regression analyses. Gaussian-smoothed firing rates (σ = 25 ms) from all units were used as predictors in a linear regression model to independently predict each axis of the tongue-tip coordinates. Thirty percent of the data were held out for testing. Because tongue-tip tracking was unavailable when the tongue was not visible in the video, firing rate traces were cropped to include only time points corresponding to valid tongue-tip measurements, ensuring frame-by-frame alignment between neural and behavioral data.

#### Spike-lick phase analysis

A lick period was defined as the inter-lick interval preceding a lick; licks lacking a preceding lick within 0.29 s were excluded. Spikes were assigned to individual licks and converted to phase values within the corresponding lick period. Spike-lick phase locking was assessed using the Rayleigh test for non-uniformity of circular data applied to the spike-phase distribution of each unit. Both p-values and vector strength were reported.

The vector strength^60^ is computed as follows: each phase was represented as a vector on the unit circle. The cosine and sine components of all vectors were averaged across observations to obtain a mean vector, and the length of this mean vector was taken as vector strength.

Statistical significance of phase non-uniformity was assessed using the Rayleigh test for circular uniformity (pycircstat implementation). The test statistic was computed from the mean vector strength and number of observations (Z = n r²), and p-values were obtained analytically using the standard exponential approximation of the Rayleigh null distribution with small-sample correction. The test evaluates unimodal deviation from a uniform circular distribution

#### Linear discriminant analysis

Linear discriminant analysis (LDA) classifiers were implemented using the LinearDiscriminantAnalysis class from scikit-learn. Default parameters were used (solver=’svd’, no shrinkage). Datasets with more than 40 relevant units (e.g MOs units for MOs dataset training) are used.

For each lick, phase values and resultant vector strength from all units were computed and concatenated to construct a matrix. The width of the matrix is 2U, since both phase value and vector strength are included for each unit; the length of the matrix is L, which is the number of licks and the above-mentioned parameters are computed for each lick. This matrix was split into train (50%) and test (50%) datasets and the split was not stratified. The train dataset was then used as input to train a linear discriminant analysis model to classify lick direction on a per-lick basis. For shuffle controls, lick direction labels were randomly permuted across trials while preserving the overall distribution of lick directions. Accuracy was computed based on the performance on test datasets.

#### Appetitive vs consummatory analysis

Only successful “go” trials in which mice maintained the same lick direction from reward-triggering licks to reward-consumption licks were included in analyses. Licks occurring within 0.7 s before reward delivery were classified as appetitive, whereas licks occurring within 0.7 s after reward delivery were classified as consummatory.

#### Noise correlation

NC between MOs unit i and MY-mot unit j was computed as the absolute Pearson correlation of their trial-to-trial mean firing rate fluctuations after subtracting the cross-trial mean. For a given time window, trial-averaged firing rates were computed by binning spikes at 5 ms resolution with 25 ms Gaussian smoothing (σ = 25 ms), then averaging across time bins within the window. NC is calculated as the pairwise correlation of the subtracted signal. Shuffle null was generated by permuting trial labels for one region (500 permutations, fixed seed) and computing NC from the permuted matrix; median shuffle NC was taken as the null per pair.

#### Spikes per lick

A lick period was defined as the inter-lick interval preceding a lick; licks lacking a preceding lick within 0.29 s were excluded. Spikes were assigned to individual licks, and the number of spikes occurring within the lick-associated time window was defined as the spike count per lick. For each unit, spike counts were averaged across all licks, including licks for which the unit did not fire.

#### Cross animal canonical correlation analysis

We analyzed medullary spiking activity from five Neuropixels recording sessions from 5 different mice performing a rewarded licking task. Spikes were aligned to the end of reward delivery and a peri-event window from −1 s to +2 s from reward was extracted (300 bins at 10 ms resolution). Only medullary premotor neurons (Allen atlas: MY-mot) that passed stability and isolation criteria were retained. Across the five datasets, the number of medullary neurons ranged from 44 to 88. Spike trains were binned, smoothed with a Gaussian kernel (σ = 20 ms). For consistency across animals, we balanced trial counts by lick direction and trial type. Within each dataset, equal numbers of “go” and “free” trials were retained for each rewarded lick direction (left, center, right). This procedure yielded 170 trials per mouse: 56 left-lick trials (44 “go”, 12 “free”), 58 center-lick trials (40 “go”, 18 “free”), and 56 right-lick trials (43 “go”, 13 “free”).

Neural data were reshaped from three-dimensional trials x time x neuron arrays into two-dimensional (trials x time) x neurons matrices. Zero-variance units were excluded, and activity was z-scored within each animal. Principal component analysis (PCA) was applied separately to each dataset, retaining 40 components per mouse. These explained between 74.8% and 98.6% of the variance in smoothed spike activity in each dataset. Multiset canonical correlation analysis (CCA, mvlearn implementation^61^) was then applied to align the five datasets. CCA in the MAXVAR formulation identifies shared latent dimensions by maximizing the variance of a consensus representation across all datasets. The model was fit with 10 canonical dimensions and ridge regularization (λ = 1 per dataset) to stabilize covariance inversions. For each mouse, CCA produced linear projections of the PCA-transformed data and latent trajectories representing activity in the shared latent space for each of the five mice.

To assess information content, multinomial logistic regression classifiers (L-BFGS solver, 5-fold cross-validation) were trained at each time bin to decode rewarded lick direction. For temporal decoding generalization tasks, classifiers were trained at each available time bin and were tested on held out fold and remaining time bins to assess decoding generalization across time. This process was completed independently for each of the 5 mice with spike data from the medullary pre-motor network. Cross animal decoding was completed by training multinomial logistic regression classifiers for each time bin on shared latent trajectories of 4 mice with 5-fold cross validation. A held-out 5^th^ mouse was used as a test set. This process was completed until all mice were represented as a test subject. Cross animal decoding as described was also completed using PCA data (unaligned) which was used as a control to assess efficacy of alignment using multiset CCA. Confusion matrices were generated by identifying maximally predicted lick direction from 0-2s time window for each trial and comparing to ground truth (rewarded lick direction). Confusion matrices were normalized across rows.

To assess if shared medullary pre-motor code generalizes to spike data from the hypoglossal nucleus, we aligned one mouse with coverage of the hypoglossal nucleus to the scores identified by aligning 5 medullary pre-motor areas. A multinomial logistic regression classifier (L-BFGS solver, 5-fold cross-validation) trained on data from the premotor areas was tested on a held out fold and on the hypoglossal nucleus.

#### Communication subspace analysis

We measured how signals are shared consistently within and across brain areas by fitting cross-validated reduced rank regression to the neural activity, following the communication subspace literature^33^. For each analysis of the communication subspace, we define a set of source units and a set of target units. When analyzing the within area currents, units are randomly assigned to either the source or the target group. Otherwise, for across-area analyses, all units recorded in each area are considered as source and target. To cross-validate across trials, we randomly assigned trials to the test or training set using 10-fold cross validation.

The training set is used to define the linear mapping between source activity and target activity via linear regression:

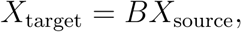

where *X_target_* is the matrix of activity *N_target_* x *T* (units x timepoints), *X_source_* is the matrix of source activity of size *N_source_* x T, and *B* is the coefficient matrix of size *N_target_* x *N_source_*. The coefficient matrix is found using least-squares regression with ridge regularization parameterized by *α*:

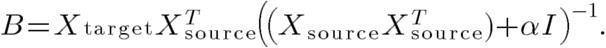

The matrix *I* corresponds to the identity matrix of size *N_source_* x *N_source_*, and the activity matrices use only trials from the training set.

The predicted target activity reads:

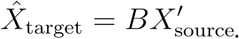

Using reduced rank regression (RRR), we then approximate *B* with a matrix *B_RRR_(r)* of lower rank *r* to determine the dimensionality of the communication subspace,

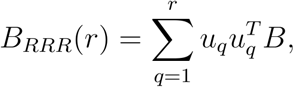

where *u_q_* corresponds to the *q*-th left singular vector of the matrix B of size *N_target_* x *1*. The predicted target using a communication subspace of dimension *r* is then computed using 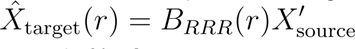. We define the optimal dimensionality *r* as the minimum rank required to reach 95% of the variance explained of the test set using *B*.

The variance explained is calculated using the *R*^2^ error between estimated and recorded activity in the target neurons in the test trials:

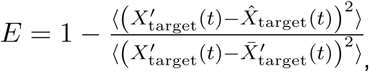

where angular brackets indicate the average over units and timepoints, 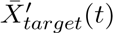 is the average firing rate across units at time *t*.

We selected neural activity 1.5 s before and 2 s after the reward delivery. The neural activity traces use the continuous signal after filtering the spike trains with a Gaussian filter of width 25ms. We defined the appetitive window from 750ms before reward until the reward, and the consummatory window from reward onset until 750ms after reward onset.

#### Recurrent neural network models

We set up a two-region recurrent neural network (RNN) whose parameters were trained to reproduce the estimated rates of recorded units in MY-mot and MOs. We focused on neural activity between 1 s before reward onset until 1.5 s after reward in each rewarded trial. We applied a Gaussian filter of 25 ms to transform spike trains into continuous rate functions. In total, we trained 10 different RNNs for each of the three sessions in which we recorded simultaneously in MY-mot and MOs.

The dynamics of the input current of the *i*-th unit in region *R* follow:

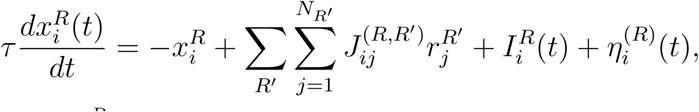

where 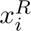 is the total current received by the neuron i in population R, 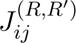 is the synaptic connection between neuron j in region R’ and neuron i in region R, 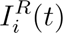 corresponds to the external input it receives, and 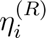 is a private Gaussian input noise term. There are three external inputs provided to the network: the reward, the tongue volume signal and the optogenetic stimulation. The reward signal is zero except for the time of reward release, and is provided to neurons in both MOs and MY-mot:

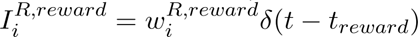. The tongue volume (TV) signal represents time-varying motor feedback and is provided to MY-mot units only: 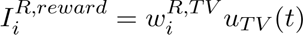 for region *R* = MY-mot. The optogenetic signal is provided to MOs units only: 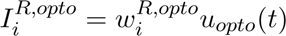 for region *R* = MOs, where *u_opto(t)_* is a signal that is one during optogenetic stimulation, zero otherwise. We refer to 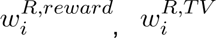, and 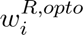 as the external input patterns.

The variable 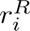 indicates the firing rate of neuron i in region R, which we are matching to the estimated rates from the data. The sum over *R’* runs over all regions in the network, in this case, MOs and MY-mot. The input-output function that transforms input current into activation is a threshold-linear, parameterized by a threshold *b* and a gain *g:* 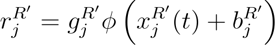. We used a membrane time constant τ of 100 ms. For numerically solving the dynamics, we used a discretization timestep of 5 ms.

We set the number of units in each region of the RNN equal to the number of recorded neurons in the corresponding dataset. A linear projection maps each unit in the data to a linear combination of the units of the same region in the RNN. The inputs received by the network correspond to the reward signal (projected to both the model MOs and MY-mot), the estimated visible volume of the tongue as a proxy for proprioceptive feedback (projected only to Med neurons), and a transient input corresponding to the initial state of all neurons at every trial. To avoid overfitting, we did not provide a quenched time-varying signal to the neurons.

The parameters that are trained during learning are the input vectors, readout vectors, recurrent connectivity weights, and single neuron gains and biases. At the beginning of training, neurons are densely connected within each region, and are sparsely connected between regions (50% sparse). The nonzero connections are randomly distributed with zero mean and variance 0.8^2^/N, where N is the total number of neurons in the network. The gains are i.i.d. Gaussian distributed with unit mean and std 0.5, the biases are i.i.d. Gaussian with zero mean and std 0.5. Discretization timestep 5 ms, membrane time constant 100 ms. To avoid instabilities, neurons receive a small amount of time-varying noise during training (std=0.001).

The loss function corresponds to the mean squared error of the RNN output and the filtered spike trains of the data. The network parameters are trained via gradient descent, using the ADAM optimizer with standard hyperparameters. We clipped gradients at a value of 1. To ensure the successful training of long time sequences, we first trained on shorter sequences of 700 ms for a fixed number of iterations, and then increased the length by 100 ms, up to reaching 2.5 s.

Once the networks are trained, we estimate the time-dependent current from every source neuron *j* to every target neuron *i* as *c_ij_(t) = J_ij_ φ(x_j_ (t))*. The set of all currents *c_ij_(t)* form a tensor of dimensions (target) neurons x (source) neurons x timepoints. To estimate the contribution of currents across areas, we averaged the norm of the currents (*c*), grouped by the region of the source neuron:

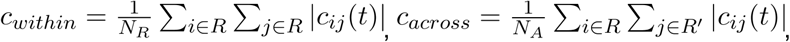

where *R* and *R’* represent the two regions (MOs and MY-mot), and *N_A_* is the number of neurons in region A. The external current is estimated as the average external input across neurons in each region.

The inhibiting stimulus provided to the RNNs that mimics optogenetic stimulation is given as a negative constant input to the RNN units in MOs. The amplitude of the stimulus is homogeneous across units and set such that the reduction in average firing rate in cortex is comparable to neural data. The stimulation period is equal to the stimulation period in the recorded data. All other external inputs in each trial are kept equal to the external inputs in the trials used during training.

## ACKNOWLEDGEMENTS

We thank Sean Quirin for building the laser collimator setup that we used for performing optogenetic stimulation during our electrophysiology experiments. We also thank Ashok Litwin-Kumar and members of the Machado lab for feedback.

## AUTHOR CONTRIBUTIONS

T.A.M. and K.D. initiated the project. T.A.M. and Y.C. performed neural recording experiments. T.A.M., Y.C., C.K., B.L., and S.B. performed behavior, optogenetics, and histology experiments. T.A.M., Y.C., M.B., M.S., and K.N., performed analyses. M.B. performed reduced rank regression analyses. M.B. and K.R. performed RNN modeling. T.A.M., Y.C., M.B., M.S. made figures. T.A.M., Y.C., M.B., M.S. and K.D. wrote the manuscript with feedback from all of the authors.

## FUNDING

T.A.M. is supported by the Howard Hughes Medical Institute, NIH award R00NS116122, and was an AP Giannini Fellow. Y.C. was supported by the Damon Runyon Cancer Research Foundation. M.B. was supported by NIH award RF1DA060772 and NSF DBI-222992. K.D. was supported by grants from the NIH, the Gatsby Foundation, and gifts from anonymous philanthropic donors.

## DECLARATION OF INTERESTS

K.D. is a founder and scientific advisor of Maplight Therapeutics and Stellaromics, and scientific advisor to RedTree and Modulight.bio. T.A.M. is a founder and board member of Numb Corp. These companies are unrelated to this work.

## DATA AND CODE AVAILABILITY

All data and code will be deposited in public repositories and made available with DOIs as of the date of publication; any additional information required to reanalyze the data is available from the lead contact upon request.

## REFERENCES

1. Allen, W.E., Kauvar, I.V., Chen, M.Z., Richman, E.B., Yang, S.J., Chan, K., Gradinaru, V., Deverman, B.E., Luo, L., and Deisseroth, K. (2017). Global Representations of Goal-Directed Behavior in Distinct Cell Types of Mouse Neocortex. Neuron 94, 891–907.e6. 10.1016/j.neuron.2017.04.017.

2. Komiyama, T., Sato, T.R., O’Connor, D.H., Zhang, Y.-X., Huber, D., Hooks, B.M., Gabitto, M., and Svoboda, K. (2010). Learning-related fine-scale specificity imaged in motor cortex circuits of behaving mice. Nature 464, 1182–1186. 10.1038/nature08897.

3. Chen, S., Liu, Y., Wang, Z.A., Colonell, J., Liu, L.D., Hou, H., Tien, N.-W., Wang, T., Harris, T., Druckmann, S., et al. (2024). Brain-wide neural activity underlying memory-guided movement. Cell 187, 676–691.e16. 10.1016/j.cell.2023.12.035.

4. Xu, D., Dong, M., Chen, Y., Delgado, A.M., Hughes, N.C., Zhang, L., and O’Connor, D.H. (2022). Cortical processing of flexible and context-dependent sensorimotor sequences. Nature 603, 464–469. 10.1038/s41586-022-04478-7.

5. Bollu, T., Ito, B.S., Whitehead, S.C., Kardon, B., Redd, J., Liu, M.H., and Goldberg, J.H. (2021). Cortex-dependent corrections as the tongue reaches for and misses targets. Nature 594, 82–87. 10.1038/s41586-021-03561-9.

6. Dragoi, T., Loschinskey, Z., Hasnain, M.A., DePasquale, B., and Economo, M.N. (2026). Dynamic engagement of the motor cortex in controlling movement. Preprint at bioRxiv, 10.64898/2026.01.13.699314 https://doi.org/10.64898/2026.01.13.699314.

7. Miri, A., Warriner, C.L., Seely, J.S., Elsayed, G.F., Cunningham, J.P., Churchland, M.M., and Jessell, T.M. (2017). Behaviorally Selective Engagement of Short-Latency Effector Pathways by Motor Cortex. Neuron. 10.1016/j.neuron.2017.06.042.

8. Hwang, E.J., Dahlen, J.E., Hu, Y.Y., Aguilar, K., Yu, B., Mukundan, M., Mitani, A., and Komiyama, T. (2019). Disengagement of motor cortex from movement control during long-term learning. Sci. Adv. 5, eaay0001. 10.1126/sciadv.aay0001.

9. Fageiry, S., Warriner, C.L., Loper, J., Pianski, L., Reardon, T., Jessell, T.M., Costa, R.M., and Miri, A. (2024). Distinct anatomical and functional corticospinal inputs innervate different spinal neuron types. Preprint at bioRxiv, 10.1101/2024.04.29.591683 https://doi.org/10.1101/2024.04.29.591683.

10. Gu, Z., Kalambogias, J., Yoshioka, S., Han, W., Li, Z., Kawasawa, Y.I., Pochareddy, S., Li, Z., Liu, F., Xu, X., et al. (2017). Control of species-dependent cortico-motoneuronal connections underlying manual dexterity. Science 357, 400–404. 10.1126/science.aan3721.

11. Stanek IV, E., Cheng, S., Takatoh, J., Han, B.X., and Wang, F. (2014). Monosynaptic premotor circuit tracing reveals neural substrates for oro-motor coordination. eLife 2014. 10.7554/eLife.02511.

12. Takatoh, J., Park, J.H., Lu, J., Li, S., Thompson, P., Han, B.-X., Zhao, S., Kleinfeld, D., Friedman, B., and Wang, F. (2021). Constructing an adult orofacial premotor atlas in Allen mouse CCF. eLife 10, e67291. 10.7554/eLife.67291.

13. Kaku, H., Gao, R., West, S., Liao, S.-M., Finkelstein, A., Kleinfeld, D., Thomas, A., Tipparaju, S.L., Svoboda, K., and Li, N. (2025). A brainstem map of orofacial rhythms. Preprint at bioRxiv, 10.1101/2025.01.27.635041 https://doi.org/10.1101/2025.01.27.635041.

14. Lindsay, N.M., Knutsen, P.M., Lozada, A.F., Gibbs, D., Karten, H.J., and Kleinfeld, D. (2019). Orofacial Movements Involve Parallel Corticobulbar Projections from Motor Cortex to Trigeminal Premotor Nuclei. Neuron 104, 765–780.e3. 10.1016/j.neuron.2019.08.032.

15. Hasnain, M.A., Han, Y., and Economo, M.N. (2025). Mapping neural subspace dynamics onto the structure of the mouse descending motor system. Preprint at bioRxiv, 10.1101/2025.10.17.682917 https://doi.org/10.1101/2025.10.17.682917.

16. Kleinfeld, D., Deschênes, M., Economo, M.N., Elbaz, M., Golomb, D., Liao, S.-M., O’Connor, D.H., and Wang, F. (2023). Low- and high-level coordination of orofacial motor actions. Curr. Opin. Neurobiol. 83, 102784. 10.1016/j.conb.2023.102784.

17. Nakamura, Y., and Katakura, N. (1995). Generation of masticatory rhythm in the brainstem. Neurosci. Res. 23, 1–19. 10.1016/0168-0102(95)90003-9.

18. Smith, J.C., Ellenberger, H.H., Ballanyi, K., Richter, D.W., and Feldman, J.L. (1991). Pre-Bötzinger complex: A brainstem region that may generate respiratory rhythm in mammals. Science 254, 726–729. 10.1126/science.1683005.

19. Takatoh, J., Prevosto, V., Thompson, P.M., Lu, J., Chung, L., Harrahill, A., Li, S., Zhao, S., He, Z., Golomb, D., et al. (2022). The whisking oscillator circuit. Nature 609, 560–568. 10.1038/s41586-022-05144-8.

20. Travers, J.B., Dinardo, L.A., and Karimnamazi, H. (1997). Motor and Premotor Mechanisms of Licking. Neurosci. Biobehav. Rev. 21, 631–647. 10.1016/S0149-7634(96)00045-0.

21. Bush, N.E., and Ramirez, J.-M. (2024). Latent neural population dynamics underlying breathing, opioid-induced respiratory depression and gasping. Nat. Neurosci. 27, 259–271. 10.1038/s41593-023-01520-3.

22. Dempsey, B., Sungeelee, S., Bokiniec, P., Chettouh, Z., Diem, S., Autran, S., Harrell, E.R., Poulet, J.F.A., Birchmeier, C., Carey, H., et al. (2021). A medullary centre for lapping in mice. Nat. Commun. 12, 6307. 10.1038/s41467-021-26275-y.

23. Ruder, L., Schina, R., Kanodia, H., Valencia-Garcia, S., Pivetta, C., and Arber, S. (2021). A functional map for diverse forelimb actions within brainstem circuitry. Nature 590, 445–450. 10.1038/s41586-020-03080-z.

24. Lindén, H., Petersen, P.C., Vestergaard, M., and Berg, R.W. (2022). Movement is governed by rotational neural dynamics in spinal motor networks. Nature 610, 526–531. 10.1038/s41586-022-05293-w.

25. Moore, J.D., Deschênes, M., Furuta, T., Huber, D., Smear, M.C., Demers, M., and Kleinfeld, D. (2013). Hierarchy of orofacial rhythms revealed through whisking and breathing. Nature 497, 205–210. 10.1038/nature12076.

26. International Brain Laboratory, Angelaki, D., Benson, B., Benson, J., Birman, D., Arlandis, J., Bonacchi, N., Bougrova, K., Bruijns, S.A., Carandini, M., et al. (2025). A brain-wide map of neural activity during complex behaviour. Nature 645, 177–191. 10.1038/s41586-025-09235-0.

27. Chen, T.W., Li, N., Daie, K., and Svoboda, K. (2017). A Map of Anticipatory Activity in Mouse Motor Cortex. Neuron 94, 866–879.e4. 10.1016/j.neuron.2017.05.005.

28. Matyas, F., Sreenivasan, V., Marbach, F., Wacongne, C., Barsy, B., Mateo, C., Chávez, M., and Petersen, C.C. (2010). Motor control by sensory cortex. Science 330, 1240–1243. 10.1126/science.1195797.

29. Wang, Q., Ding, S.-L., Li, Y., Royall, J., Feng, D., Lesnar, P., Graddis, N., Naeemi, M., Facer, B., Ho, A., et al. (2020). The Allen Mouse Brain Common Coordinate Framework: A 3D Reference Atlas. Cell 181, 936–953.e20. 10.1016/j.cell.2020.04.007.

30. Deschênes, M., Takatoh, J., Kurnikova, A., Moore, J.D., Demers, M., Bouvier, J., Lee, D., and Kleinfeld, D. (2016). Control of orofacial motor actions by vibrissa motor cortex. Nature 538, 301–305. 10.1038/nature19766.

31. Safaie, M., Chang, J.C., Park, J., Miller, L.E., Dudman, J.T., Perich, M.G., and Gallego, J.A. (2023). Preserved neural dynamics across animals performing similar behaviour. Nature 623, 765–771. 10.1038/s41586-023-06714-0.

32. Carroll, J.D. (1968). Generalization of Canonical Correlation Analysis to Three of More Sets of Variables. 10.1037/e473742008-115.

33. Semedo, J.D., Zandvakili, A., Machens, C.K., Yu, B.M., and Kohn, A. (2019). Cortical Areas Interact through a Communication Subspace. Neuron 102, 249–259.e4. 10.1016/j.neuron.2019.01.026.

34. Perich, M.G., Arlt, C., Soares, S., Zhou, S., Beiran, M., Andalman, A.S., Benster, T., Young, M.E., Mosher, C.P., Minxha, J., et al. (2026). Inferring brain-wide interactions using data-constrained recurrent neural network models. Neuron, S0896-6273(26)00571-4. 10.1016/j.neuron.2026.07.016.

35. Gerstner, W. (2000). Population Dynamics of Spiking Neurons: Fast Transients, Asynchronous States, and Locking. Neural Comput. 12, 43–89. 10.1162/089976600300015899.

36. O’Shea, D.J., Duncker, L., Goo, W., Sun, X., Vyas, S., Trautmann, E.M., Diester, I., Ramakrishnan, C., Deisseroth, K., Sahani, M., et al. (2022). Direct neural perturbations reveal a dynamical mechanism for robust computation. Preprint at bioRxiv, 10.1101/2022.12.16.520768 https://doi.org/10.1101/2022.12.16.520768.

37. Hwang, E.J., Dahlen, J.E., Mukundan, M., and Komiyama, T. (2021). Disengagement of Motor Cortex during Long-Term Learning Tracks the Performance Level of Learned Movements. J. Neurosci. 41, 7029–7047. 10.1523/JNEUROSCI.3049-20.2021.

38. Koh, N., Ma, Z., Sarup, A., Kristl, A.C., Agrios, M., Young, M., and Miri, A. (2025). Selective direct influence of motor cortex on limb muscle activity during naturalistic climbing in mice. Nat. Neurosci. 28, 2537–2549. 10.1038/s41593-025-02093-z.

39. Fries, P. (2015). Rhythms for Cognition: Communication through Coherence. Neuron 88, 220–235. 10.1016/j.neuron.2015.09.034.

40. Karalis, N., and Sirota, A. (2022). Breathing coordinates cortico-hippocampal dynamics in mice during offline states. Nat. Commun. 13, 467. 10.1038/s41467-022-28090-5.

41. Hsueh, B., Chen, R., Jo, Y., Tang, D., Raffiee, M., Kim, Y.S., Inoue, M., Randles, S., Ramakrishnan, C., Patel, S., et al. (2023). Cardiogenic control of affective behavioural state. Nature 615, 292–299. 10.1038/s41586-023-05748-8.

42. Marder, E., and Bucher, D. (2001). Central pattern generators and the control of rhythmic movements. Curr. Biol. 11, R986–R996. 10.1016/S0960-9822(01)00581-4.

43. Komi, S., Kaur, J., Winther, A., Bonfils, M.C.A., Houser, G.A., Sørensen, R.J.F., Li, G., Sobriel, K., and Berg, R.W. (2025). Neural manifolds that orchestrate walking and stopping. Preprint at bioRxiv, 10.1101/2025.11.08.687367 https://doi.org/10.1101/2025.11.08.687367.

44. Economo, M.N., Viswanathan, S., Tasic, B., Bas, E., Winnubst, J., Menon, V., Graybuck, L.T., Nguyen, T.N., Smith, K.A., Yao, Z., et al. (2018). Distinct descending motor cortex pathways and their roles in movement. Nature 563, 79–84. 10.1038/s41586-018-0642-9.

45. McElvain, L.E., Friedman, B., Karten, H.J., Svoboda, K., Wang, F., Deschênes, M., and Kleinfeld, D. (2018). Circuits in the Rodent Brainstem that Control Whisking in Concert with Other Orofacial Motor Actions. Neuroscience 368, 152–170. 10.1016/j.neuroscience.2017.08.034.

46. Perich, M.G., and Rajan, K. (2020). Rethinking brain-wide interactions through multi-region ‘network of networks’ models. Curr. Opin. Neurobiol. 65, 146–151. 10.1016/j.conb.2020.11.003.

47. Machado, T.A., Kauvar, I.V., and Deisseroth, K. (2022). Multiregion neuronal activity: the forest and the trees. Nat. Rev. Neurosci. 23, 683–704. 10.1038/s41583-022-00634-0.

48. Das, A., and Fiete, I.R. (2020). Systematic errors in connectivity inferred from activity in strongly recurrent networks. Nat. Neurosci. 23, 1286–1296. 10.1038/s41593-020-0699-2.

49. Beiran, M., and Litwin-Kumar, A. (2025). Prediction of neural activity in connectome-constrained recurrent networks. Nat. Neurosci. 28, 2561–2574. 10.1038/s41593-025-02080-4.

50. Huang, A., Singh, S.H., Martinelli, F., and Rajan, K. (2025). Measuring and Controlling Solution Degeneracy across Task-Trained Recurrent Neural Networks. Preprint at arXiv, 10.48550/arXiv.2410.03972 https://doi.org/10.48550/arXiv.2410.03972.

51. Kauvar, I.V., Machado, T.A., Yuen, E., Kochalka, J., Choi, M., Allen, W.E., Wetzstein, G., and Deisseroth, K. (2020). Cortical Observation by Synchronous Multifocal Optical Sampling Reveals Widespread Population Encoding of Actions. Neuron 107, 351–367.e19. 10.1016/j.neuron.2020.04.023.

52. Richman, E.B., Ticea, N., Allen, W.E., Deisseroth, K., and Luo, L. (2023). Neural landscape diffusion resolves conflicts between needs across time. Nature 623, 571–579. 10.1038/s41586-023-06715-z.

53. Allen, W.E., Chen, M.Z., Pichamoorthy, N., Tien, R.H., Pachitariu, M., Luo, L., and Deisseroth, K. (2019). Thirst regulates motivated behavior through modulation of brainwide neural population dynamics. Science 364, eaav3932. 10.1126/science.aav3932.

54. Jun, J.J., Steinmetz, N.A., Siegle, J.H., Denman, D.J., Bauza, M., Barbarits, B., Lee, A.K., Anastassiou, C.A., Andrei, A., Aydin, Ç., et al. (2017). Fully integrated silicon probes for high-density recording of neural activity. Nature 551, 232–236. 10.1038/nature24636.

55. Steinmetz, N.A., Aydin, C., Lebedeva, A., Okun, M., Pachitariu, M., Bauza, M., Beau, M., Bhagat, J., Böhm, C., Broux, M., et al. (2021). Neuropixels 2.0: A miniaturized high-density probe for stable, long-term brain recordings. Science 372, eabf4588. 10.1126/science.abf4588.

56. Fiáth, R., Márton, A.L., Mátyás, F., Pinke, D., Márton, G., Tóth, K., and Ulbert, I. (2019). Slow insertion of silicon probes improves the quality of acute neuronal recordings. Sci. Rep. 9, 111. 10.1038/s41598-018-36816-z.

57. Tyson, A.L., Rousseau, C.V., and Margrie, T.W. (2020). brainreg: automated 3D brain registration with support for multiple species and atlases. (Zenodo). 10.5281/zenodo.3991718 https://doi.org/10.5281/zenodo.3991718.

58. Pachitariu, M., Sridhar, S., Pennington, J., and Stringer, C. (2024). Spike sorting with Kilosort4. Nat. Methods 21, 914–921. 10.1038/s41592-024-02232-7.

59. Siegle, J.H., Jia, X., Durand, S., Gale, S., Bennett, C., Graddis, N., Heller, G., Ramirez, T.K., Choi, H., Luviano, J.A., et al. (2021). Survey of spiking in the mouse visual system reveals functional hierarchy. Nature 592, 86–92. 10.1038/s41586-020-03171-x.

60. Goldberg, J.M., and Brown, P.B. (1969). Response of binaural neurons of dog superior olivary complex to dichotic tonal stimuli: some physiological mechanisms of sound localization. J. Neurophysiol. 32, 613–636. 10.1152/jn.1969.32.4.613.

61. Perry, R., Mischler, G., Guo, R., Lee, T., Chang, A., Koul, A., Franz, C., Richard, H., Carmichael, I., Ablin, P., et al. (2021). mvlearn: Multiview Machine Learning in Python. Preprint at arXiv, 10.48550/arXiv.2005.11890 https://doi.org/10.48550/arXiv.2005.11890.

